# A mechanistic model towards ecological inclusion for a better plant growth understanding

**DOI:** 10.1101/2022.12.09.519755

**Authors:** Lucia Nasti, Emanuela Del Dottore, Fabio Tedone, Michele Palladino, Barbara Mazzolai, Pierangelo Marcati

## Abstract

Plants growth is a complex and delicate balance among different factors involving environmental and physiological conditions. In this context, we propose a mechanistic model that considers the main internal processes of plant growth and reproduces a wide range of plant behaviors observed experimentally. In particular, we describe the model plant of *Arabidopsis thaliana* in a realistic environment with a day-and-night cycle, which considers different inputs as light, water, phosphorus, nitrogen, starch and sucrose. In addition, we propose a new function that describes the affinity between the plant and a specific nutrient, and a novel feedback signal to model how plants have the remarkable capacity to distribute resources among their organs. The result is an efficient tool applicable in ecological and agricultural studies, which can estimate several parameters, simulate several soil conditions and analyze how limiting or toxic resources can affect the plant development. We improve our understanding of plant adaptive strategies, reproducing results in line with experiments.

## 1. Introduction

Plants are living organisms coordinating a complex network of internal, e.g., nutrient concentration, and external signals, e.g., light and soil resources. As a result, plant growth is the consequence of a delicate balance among different factors involving environmental and physiological conditions [1].

To investigate on their adaptive strategies in multiple situations, we focus on two specific processes that depend on soil characteristics: the nutrient uptake and the allocation of resources among roots and shoot.

Generally, we assume that nutrient uptake rates follow Michaelis-Menten kinetics [2], whose parameters change according to soil resources and plants’ internal status. To model resources allocation [3], e.g., sucrose or starch,, we can use the notion of *Sink priority* [4], by which resources move from a region where are generated or stocked (sources) to a region where are consumed (sink). All plant tissues, which passively receive the sucrose and are photosynthetically inactive, are represented as sinks. The priority of each sink is dynamically determined by the plant according to external stimuli. For example, when the soil is poor in nutrients, the roots (representing the sink) will require more energy to uptake the nutrients and, consequently, they will need more sucrose.

Analyzing these mechanisms is complicated because they are profoundly interconnected and characterized by multiple adaptive responses that make it difficult to isolate individual functions, e.g., photosynthesis, nutrients inter-relation, energy demand for growth. In this context, a possible approach is to build a realistic model, crucial to achieving an integrated view of all biological phenomena involved in plant development.

Mechanistic models typically involve physically interpretable parameters, allowing more in-depth insights into system performance and better predictions of its behaviour [5]. Even if these kinds of models require a priori information on the system and often need more time and resources for validation, it is possible to incorporate empirical concepts and biological assumptions to reproduce the plant complexity as faithfully as possible [6].

In literature, many works exploit mathematical models to focus on specific biological processes. For example, the work in [7] analyses the role of nitrogen in the photosynthesis. In [8], the authors model the function of nitrogen in plant respiration. In [9], they focus on the allocation of sucrose among different organs, and in [6], the authors propose the first combination of photosynthesis, growth, and phosphorus uptake. Finally, in [10], the authors examine starch and sucrose production and how they are related to the circadian clock.

In this work, we propose a mechanistic model that considers the main internal processes of plant growth and reproduces a wide range of plant behaviors observed experimentally. We describe the growth of the model plant *Arabidopsis thaliana* in a realistic environment with a day-and-night cycle. We include light, water, phosphorus, and nitrogen as resources and inputs of the system affecting leaves and roots biomass. Starch and sucrose are both involved: in particular, we model the complex dynamics of the carbohydrate reserves essential to emulate the switch between light and dark periods. We also combine the costs for respiration, transport, growth, and nutrient uptake.

In this context, we provide the main novelty of our model. To describe the nutrient uptake, we propose two new functions that emulate an internal signal expressing the affinity between the plant and a specific nutrient (nitrogen or phosphorus). These two functions (*a_n_*(*t*) and *a_p_*(*t*)) simulate - simply and efficiently - how plants interact with soil nutrients, considering the plant status, such as the photosynthesis activity and the sucrose consumption, and not only the nutrient concentration in soil. This approach allows us to define precisely the uptake process, combining different aspects of plant behaviour.

In addition, we introduce a feedback signal to model how plants have the remarkable capability to coordinate the growth of their organs, distributing the resources among them. Then, we can balance the nutrient demand for the most limiting resource and describe the sucrose allocation between leaves and roots. These new formalisms allow us to simulate several real scenarios, such as plant starvation, photosynthesis boost, absence of balance among nutrients (i.e., *stoichiometry ratio*).

To calibrate the parameters of our functions, we mainly use the works done in [11, 12, 13]. Then, we proceed by validating this model with several independent experimental data sets. In addition, we compare our results with the results obtained by the model described in [7] to show the accuracy and the predictivity of our work.

The result is an efficient tool applicable in ecological and agricultural studies, which can estimate several parameters, such as biomass and sucrose production, rate of nutrient uptake, and the relative growth rate. Further, it can simulate several soil conditions, analyzing how limiting or toxic resources can affect plant development. By integrating different resources and processes in a single comprehensive model, we improve our understanding of physiological processes and plant adapting strategies, reproducing results in line with experimental observations. With our approach, we aim at developing a model that can better fit plant growth dynamics with its ecological niche, by including interlaced functions that integrate the effects of environmental conditions in the internal state of the plant.

Indeed, understanding the principles of plant growth is fundamental in many research fields, such as ecology, agriculture, and genetics, among others. In ecological studies, the authors investigate the role of natural and mechanical factors, such as light and wind, on the phenotypic variation of the plant during development. In [14], for instance, they examine the external causes (i.e., climate change, atmospheric CO2 concentrations) that produce abrupt changes in plants during growth, which lead to the evolution of some plants’ specific features.

Plant genetics focus on growth and development to predict phenotypic traits from new genotypes under untested environmental conditions. Then, it is possible to build a mechanistic model to simulate breeding strategies to improve target traits [15].

A detailed model of plants’ functioning can support agriculture and forestry. In agriculture, a central interest is to maximize the effects of fertilizer and nutrients to improve production, which can be boosted by knowing how plants take up nitrogen and other elements. In general, one of the typical coefficients to examine agriculture production is the Nutrient Use Efficiency parameter (NUE), which estimates how specific supplements added in soils can affect crop yields [16, 17]. However, to obtain a more accurate estimation, we need to use the NUE value in combination with other factors [18], such as the rate of the nitrogen productivity, the nutrient concentrations, the photosynthesis rate, and others, thus complicating the analysis.

Another novel field of appliation for this study is in plant-inspired robotics. In fact, deepening the knowledge on plant functioning, including metabolism and architecture evolution,help to realize biomimetic autonomous systems that on one side can better adapt in challenging and dynamic environments and on the other hand by better imitating plant behaviors can be adopted as platforms to support the study of the biological model [19].

This work is organized as follows. In Section 2, we provide a clear illustration of the assumptions and hypotheses that we consider in our model. In Section 3, we describe how we integrate the functions and estimate the parameters, while in Section 4, we compare our work to the work done by Franklin et al. [7]. In Section 5, we outline our results and illustrate how this mechanistic model is a valuable tool to analyze complex plant behaviors. Finally, in Section 6, we draw our conclusions and discuss future work.

## 2. Description of the model

### 2.1. General hypothesis

This model consists of mechanistics and biological principles necessary to understand plant growth. Here, we provide a clear illustration of the assumptions and hypotheses that we consider in our model, which is represented schematically in Figure 1. In Table 1, we collect all the model’s functions, which are largely detailed in the Supplementary Material as well.

**Figure 1:**
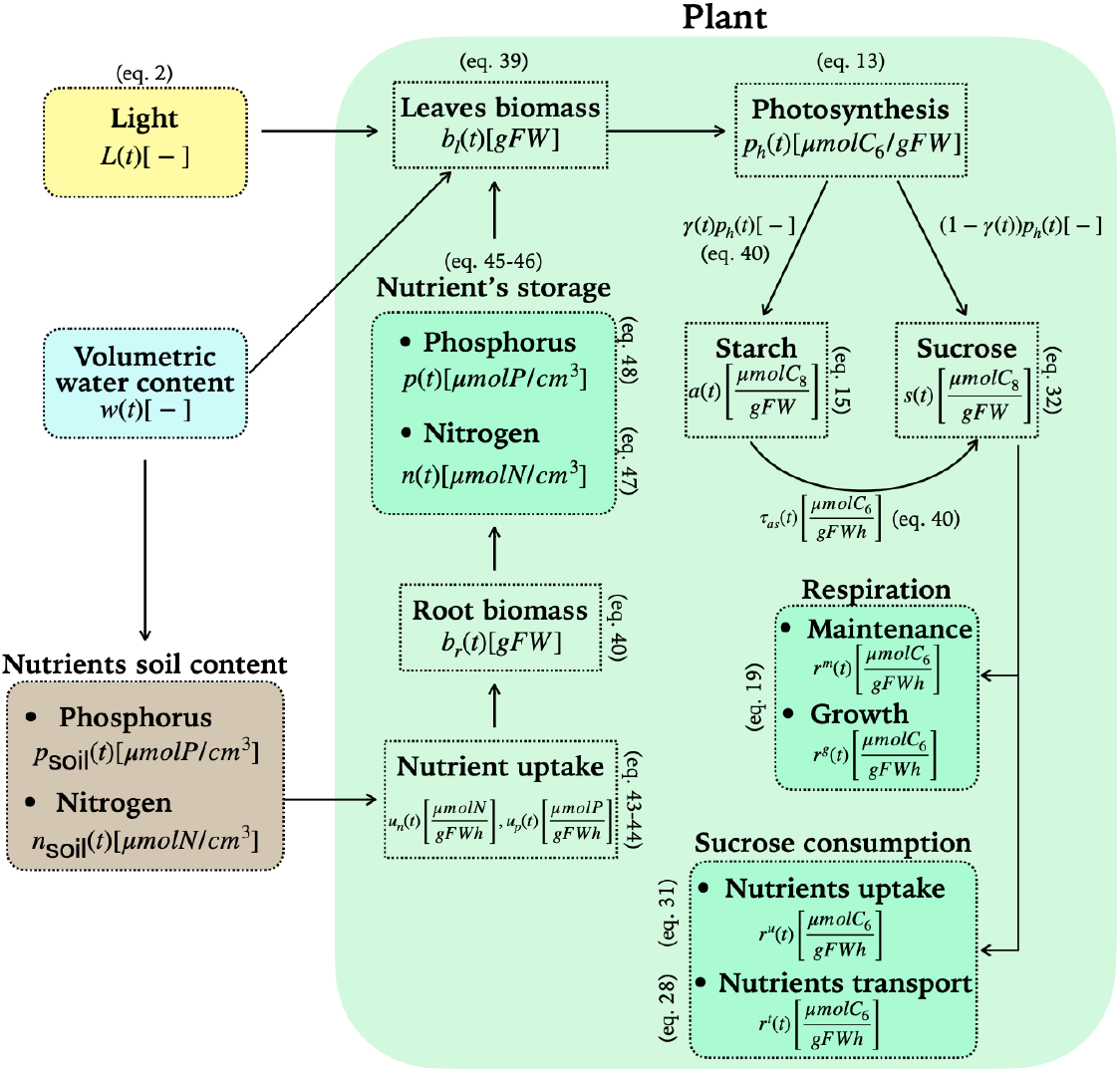
Schematic representation of the plant processes and the external resources involved in our model.

**Table 1:** Functions constituting the mathematical model.

| Functions | Description |
| --- | --- |
| $p_h(t) = p_h^{\max} L(t) \bar{w}(t) \min(\bar{n}(t), \bar{p}(t)), p_{\text{tox}}(t, p_{\text{soil}}(t)) C(t, a(t)) \frac{b_l(t)}{b_l(t) + \theta_{bl}}$ | Photosynthesis |
| $\frac{d\gamma(t)}{dt} = L(t) (-\gamma(t) \lambda_{sdr} \frac{s^{\min}}{s(t) + s^{\min}} + (1 - \gamma(t)) \lambda_{sdi} \frac{s(t)}{s(t) + s^{\max}}) + (1 - L(t)) (1 - \gamma(t)) \lambda_{sni} \frac{s^{\min}}{s^{\min} + s(t)}$ | Starch accumulation |
| $\frac{ds(t)}{dt} = (1 - \gamma(t)) p_h(t) + \tau_{as}(t) - r^u(t) - r^t(t) - r^m(t) - r^g(t)$ | Sucrose dynamics |
| $\frac{db_l}{dt} = \lambda_{sb} (1 - f_r(t) n_{tox}(t)) r^g(t) b_l(t) - \theta_{w2}(t) \theta_{ld} b_l(t) - \theta_{lc} b_l^2(t)$ | Leaves growth |
| $\frac{db_r}{dt} = \lambda_{sb} f_r(t) n_{tox}(t) r^g(t) b_l(t) - \theta_{rd} b_r(t) - \theta_{rc} b_r^2(t)$ | Root growth |
| $u_n(t) = u_{nMM}(I_n^{\max}, k_n^{\max}) a_n(t) u_n^{\text{sat}}(t) \frac{b_r(t)}{b_l(t) + b_r(t)}$ | Nitrogen uptake |
| $u_p(t) = u_{pMM}(I_p^{\max}, k_p^{\max}) a_p(t) u_p^{\text{sat}}(t) \frac{b_r(t)}{b_l(t) + b_r(t)}$ | Phosphorus uptake |
| $r^m(t) = \bar{r}_m(t) s(t)$ | Maintenance |
| $r^g(t) = \bar{r}_g(t) s(t)$ | Growth |
| $\frac{df_r(t)}{dt} = p_{lux}(t) (1 - f_r(t)) (a_n(t) f_n(t) + (1 - f_n(t)) a_p(t)) - f_r(t) \left( \frac{n(t)(1 - f_n(t))}{n(t) + n^{\min}(t)} + \frac{p(t) f_n(t)}{p(t) + p^{\min}(t)} + \frac{s^{\min}}{s(t) + s^{\min}} \right)$ | Root's priority |

Four main stages characterize the life cycle of most plants: germination of the seeds, vegetative phase, flowering and maturation of fruits, and senescence. In our work, we refer to *Arabidopsis thaliana* since it is a widely study model plant with largely available information. We focused on its vegetative state since it is the most active period for plant development. As described in [20], at this stage, the plant uses its energy and biomass to grow and to accumulate resources needed for the following flowering and reproductive phases. In addition, we assume that the model has two compartments: the above-ground leaf biomass, relevant for the photosynthesis, and the below-ground root biomass, mainly devoted to the nutrient uptake. Other essential assumptions are:

- Water availability limits plant development because it increases the nutrient uptake;
- Light *L*(*t*)[-] influences all the biological processes, as photosynthesis;
- As nutrients, we include nitrogen (*N*), a fundamental constituent of protein and chlorophyll, promoting root growth and nutrients uptake, and phosphorus (*P*), which stimulates root lengthening and supports the energy carriers within the plant;
- As established in [21, 22], the nutrient uptake rate follows the Michaelis-Menten kinetics. Indeed, the nutrient transportation is similar to the reaction enzyme-substrate [2];
- In order to represent natural conditions in the context of plant growth, we include many phenomena as the death of tissues, the self-shading effect (due to an overproduction of leaves), and the self-competition in roots limiting plant growth;
- We measure the sugar (sucrose or starch) in *μmolC*_6_, while the biomass is measured in grams of fresh weight ([*gFW*]). We estimate the conversion from fresh weight ([*gFW*]) to dried weight ([*gDW*]) as *FW* = 12.5*DW*, as done in [11].

### 2.2. Plant processes and resources involved in the model

#### 2.2.1. Photosyntesis

The photosynthesis is an integral part of plant metabolism used to produce carbohydrates. Leaf biomass, water, light, and nutrient concentration affect this process [23]. Thus, to model it, we take into account all these factors.

First of all, we use a function (*p_h_*(*t*)[*pmolC*_6_/*gFW* · *h*]) that predicts the rate of sugar production for each gram of fresh photosynthetically active biomass, namely the leaf biomass *b_l_*:

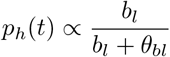

where *θ_bl_* is a parameter that has to be estimated. We assume that if there are no leaves, we have *b_l_* = 0, and then *p_h_* = 0. Otherwise, the photosynthesis rate increases with increasing leaf number.

We define light *L*(*t*) as:

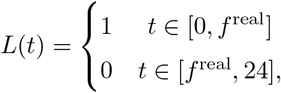

where *f*^real^ represents the environmental or real photoperiod, namely the number of hours of light during the day, when photosynthesis is actually effective. Usually, *f*^real^ corresponds to the plant perceived photoperiod *f* (*t*), then we can write *f*^real^ = *f* (*t*). If it suddenly changes, the plant needs time to adapt its metabolism to the new day length [24]. Then, we assume that *f*^real^ and *f* (*t*) will match again after 24 hours.

We add to the model the influence of water amount (*w_soil_*(*t*)[-]) on photosynthesis using the following saturation function:

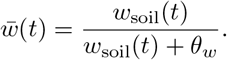

This function reproduces when insufficient water intake, i.e., *w_soil_* → 0, reduces photosynthesis, which - in normal conditions - can reach its maximal level 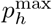.

In addition, we model how nitrogen and phosphorus change the rate of carbon production, as explained in [25, 26]. In particular, toxic levels of phosphorus in the soil *p_soil_*[*μmolP*/*cm*^3^] can drop the photosynthesis production up to 50% [27].

The reduction of photosynthesis rate may depend on the starch concentration, which is one of its products. Indeed, plants collect starch to avoid starvation during the night. However, when the starch storage is sufficient, plants can limit carbohydrates production, modifying the stomatal activity [28]. To describe this phenomenon, we implement the *photosynthetic control function C*(*t, a*(*t*)), where *a*(*t*) represents the starch content at time *t*.

#### 2.2.2. Starch and sucrose

Sucrose and starch are photosynthetic products necessary to sustain diurnal and nocturnal metabolism [29]. Plants balance their production and storage: when the sucrose consumption increases, they lower the accumulation of starch that, instead, increases during nightly starvation [6, 30, 31]. In this context, we implement a dynamic function, the *starch accumulation signal γ*(*t*) ∈ [0,1][-], which simulates a trigger of starch production in the case of rising sucrose levels. The same equation inhibits starch accumulation during daily sucrose starvation, modeled by estimating a minimum sucrose threshold (*s*^min^) that the plant needs to preserve [32]. To describe the starch degradation and conversion in sucrose during the night, we design another function, *τ*_as_(*t*), that relies on the photoperiod.

Photoperiod, indeed, scans the plant’s activity: while plants use starch in the night, during the day, they absorb sucrose to sustain living tissues (maintenance respiration) and grow up (growth respiration). Then, we model the sucrose dynamics using the *s*(*t*) function, considering all the metabolic processes producing sucrose, such as photosynthesis, and consuming it, as nutrient uptake, maintenance, and growth respiration.

The sucrose concentration is particularly relevant in the model because, as shown in [33], we assume that maintenance *r*^m^(*t*) and growth *r*^g^(*t*) are proportional to *s*(*t*):

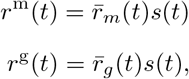

where 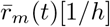 and 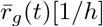 are proportionality functions representing the frequencies of loading sucrose into the phloem during the respiration. We extend the definition of 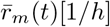, including two terms: a frequency term for the loading of sucrose 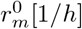, and a frequency parameter *θ_s_* [1/*h*] that simulates the losses of sucrose during transportation because of the porous structure of the phloem as described in [34]. Therefore, we specify 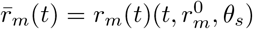. To characterize the growth respiration, we write 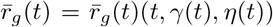. Here, we use two functions: *γ*(*t*) expresses the effect of nightly starvation on growth, while *η*(*t*) the deficiency of nutrients in the soil. In particular, this aspect induces a higher second metabolism and stronger defense stimulus [35].

Sucrose is divided between roots (denoted as *f_r_* (*t*) ∈ [0,1][–]) and leaves (1 – *f_r_* (*t*)) [3, 36, 18]. In general, the roots’ activity affects the amount of sucrose invested in their growth. For instance, if the soil is rich in nutrients, they need less energy because the assimilation of minerals is easier. A counterintuitive behavior is verified when the ground contains toxic phosphorus levels [37]. In this case, the plant intakes more phosphorus than necessary, namely *luxury uptake*, which inhibits growth. Other conditions can cause growth inhibition. An example is when plants prioritize leaves development to benefit from a photosynthesis booster.

#### 2.2.3. Nutrient uptake

Nutrients govern many metabolic activities like photosynthesis [8], sucrose assimilation in tissues [7], starch storage [38], and respiration [6].

To describe the uptake rate of nutrients, we can use the Michaelis-Menten kinetics as shown in [2]. The formulas describing the rates for nitrogen and phosphorus are the following:

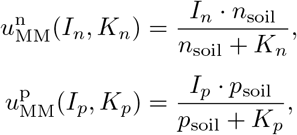

where *I_n_, k_n_, I_p_*, and *k_p_* are Michaelis-Menten parameters, whose value depends on the soil conditions, which implies a new estimation every time they change. To overcome this limitation and simplify the modelization, we assume the parameters as fixed and set to their highest value, and we design two internal plant’s affinity signals, namely *a_n_*(*t*) and *a_p_*(*t*) respectively for nitrogen and phosphorus, to limit the uptake rate.

This contribution represents the main novelty of our model. Indeed, the signals *a_n_*(*t*) and *a_p_*(*t*) ∈ [0,1] allow us to simulate the affinity between the plant and the specific nutrient, describing different possible scenarios.

When *a_n_*(*t*) = 1 and *a_p_*(*t*) = 1, the rates are at their highest level, coinciding with 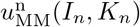 and 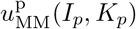. Otherwise, with *a_n_*(*t*) = 0 and *a_p_*(*t*) = 0, the rates are turned off even if the nutrients are abundant in the soil.

We assume that the signals increase when:

- the plants are consuming more nutrients;
- the photosynthetic activity increases;
- there is no balance between nitrogen and phosphorus amounts. In this case, we talk about the ratio among the plant’s nutrient contents, known as *stoichiometry ratio*.

The signals decrease when:

- the plant already stored part of the nutrients;
- the sucrose consumption required to store the nutrients is limiting to support the other biological processes.

In this context, the challenge is to define a meaningful threshold to specify the concentration of nutrients. To solve this problem, we estimate two lower bounds, *n*^min^(*t*) and *p*^min^(*t*), and two upper bounds, *n*^max^(*t*) and *p*^max^(*t*). The lower bounds estimate the concentration of nutrients necessary to maintain the metabolism in the worst conditions when the consumption of sucrose and starch is maximum for 24h. Instead, the upper bounds simulate the highest concentration of nutrients that the plants can store. This quantity depends on the number of days (*D*) the plant could sustain its metabolism in a sudden total absence of nutrients. As described in [8], for herbaceous plants, like the *Arabidopsis thaliana*, the number *D* is four days.

In addition, we analyze and include conditions and effects of toxicity and deficiency levels of nutrients in the soil because they can modify the plant’s development. For example, as pointed out in [35], the secondary metabolism of the plant overcomes its growth when the soil is poor on nutrients. In particular, as shown in [39], we model how a toxic level of phosphorus can induce in the plant the identical consequences of an iron deficiency.

## 3. Model implementation and parameter estimation

We use the numerical solver *ODE45* of Matlab to integrate the model differential equations. In Table 4 of Supplementary Material, we list the initial conditions. The model contains 50 parameters. Among these, we collect 21 from literature, 11 by fitting the biological results reported in [11], 4 considering the work in [12], and 6 from [13]. In addition, we specify 8 parameters arbitrarily because the recalled experiments do not observe their specific effects.

Unless otherwise noted, we fix phosphorus and nitrogen to satisfy the stoichiometry ratio in no-limiting conditions for plant growth. The control signals are set to 0.5 to start from the equilibrium.

Going into details about the parameter estimation, we mainly use the works done in [11, 12, 13].

Using the data in [11], we estimate the effect of toxic levels of phosphorus in the soil. Here, the authors grew the plant for five days at 16h photoperiod, and then, after seven days, they changed the phosphorus concentration testing different treatments. They observe the impact of this element on the root length, which can show a decrease of up to 50%. We collect the found parameters in Table 3 in Supp. Material.

In [12], the authors grew *Arabidopsis* under five different photoperiods (4, 6, 8, 12, and 18 hours of light) for 29 days. From their results, we extract the parameters that we need to calibrate the functions, such as the rate of the sucrose conversion in new biomass (*λ_sb_*) and the frequency parameters in the dynamics of the starch accumulation signal (*γ*(*t*)), as reported in Table 4 in Supp. Material.

Finally, we consider the work in [13] to study the consequences of poor or toxic levels of nitrogen on plant growth. The authors grew *Arabidopsis* for 35 days in 8h photoperiod in no-limiting phosphorus soil conditions. In particular, from this work, among the other parameters that we collect in Table 4 of Supp. Material, we estimate *δ_npd_* to model the minimum growth stimulus reduction in nutrients deficiency.

## 4. Comparison with the model of Franklin et al. (2017)

Many works and experiments have observed the importance of nitrogen for plant nutrition [40, 41, 42]. In particular, the work done by Franklin et al. [7] analyzes how nitrogen may significantly influence plant shoot and root growth by multiple laboratory experiments involving different species (Arabidopsis, poplar, pine, and spruce). To focus on the effects of nitrogen availability, they propose a mathematical model, which examines only the essential biological processes involving carbon assimilation and cost. The model consists of two differential equations describing biomass growth and nitrogen assimilation.

We implement their model to compare the performance of our works on several experimental results. We aim to show that our model stands out for accuracy and predictivity.

Indeed, at the structural level, Franklin’s and our model follow - in part - the same assumptions: they consider the plant as divided into two parts (shoot and root compartments), and growth and maintenance respiration are proportional to nitrogen concentration. On the other hand, they do not include many external inputs, such as water, or other needed nutrients, such as phosphorus, and they do not model the starch effects on plant development.

### 4.1. Comparison of the two models

For an initial comparison of the two models, we use the experimental data of the paper of Sulpice et al. [11]. Here, the authors want to study how disparate photoperiods can affect plants growth. As a metric, they use the Relative Growth Rate (RGR): the rate of increase of total weight (dry or fresh) per plant, *W*, expressed per unit of *W*.

In Table 2, we examine in contrast the RGR of our model, the model of Franklin et al. [7], and the data produced by the experiments conducted in [11]. As we can notice, our results are in agreement with the observations of [11], and this is widely attributable to the examination of the sucrose and starch dynamics in both works.

**Table 2:** The mean values of RGR. With respect to the experimental data of Sulpice et al. [11], we compare the mean value of RGR of Franklin et al.’s [7] and our models.

|  | Photoperiod |  |  |  |  |
| --- | --- | --- | --- | --- | --- |
| Source | 4h | 6h | 8h | 12h | 18h |
| Sulpice et al. (Data) | 0.068 | 0.1135 | 0.1708 | 0.26 | 0.3065 |
| Our model | 0.068 | 0.1135 | 0.1708 | 0.2598 | 0.3065 |
| Franklin et al. (Model) | 0.048 | 0.074 | 0.099 | 0.1501 | 0.2259 |

In [12], the authors study the effect of toxic nutrient concentration to understand how the absorption of phosphorus can influence plants development. They provide various treatments experimentation, such as the effects of total absence of phosphorus and the toxic soil concentration (i.e., *2μmolP/cm^3^*). Since the model in [7] does not include the phosphorus uptake, in Table 3, we compare the estimation of the root biomass, considering the case in which the phosphorus is absent. As reported in the example before, also, in this case, our model agrees with the experimental results.

**Table 3:** The root biomass estimation. With respect to the observations produced by the work of Shulka et al. [12], we compare the root biomass estimation of Franklin et al.’s [7] and our models.

| Source | Root Biomass Estimation |
| --- | --- |
| Shukla et al. (Data) | 24.8$\mu gFW$ |
| Our model | 25.14$\mu gFW$ |
| Franklin et al. (Model) | 11.5$\mu gFW$ |

By these two comparisons, we aim to show the applicability extent of our work: having a comprehensive model, which includes many internal and external inputs, guarantees a better understanding of the plant dynamics in line with the data.

## 5. Results

Our mechanistic model is a valuable tool to analyze complex plant behaviors: we can estimate sucrose and biomass productions and understand nutrient dynamics. Furthermore, we can improve our knowledge about physiological processes and plant adapting strategies.

### 5.1. Starch and sucrose dynamics

Starch and sucrose are two essential components of plant development, which have a prominent role in biotechnologies. For instance, the production of biofuels, like bioethanol, requires starch fermentation [43], which is then indispensable to estimate: to improve the biofuels industry is crucial to understand the environmental factors affecting plants growth and, in particular, starch dynamics.

We show that our model estimates starch and sucrose dynamics in agreement with the observations presented in literature [44]. Here, the authors investigate how starch responds to changes in photoperiod and irradiance. To do that, they grow plants in a short (6h) and a longer (12h) photoperiod using three light intensities at each photoperiod. As reported in Table 1 in [44], they observe that a lower intensity light (160*μmol·m*^-2^ · *s*^-1^) produces a decrement of 14.4-24.4% in starch concentration comparing short and long photoperiod. Instead, a higher light intensity (320*μmol*· *m*^-2^ ·*s*^-1^) causes a diminution of 10.9-31.4% of starch.

To compare these results with the predictions provided by our model, we set the light intensity to 160*μmol*·*m*^-2^ ·s^-1^, and simulate the growth of the plant for 30 days, measuring the concentration every day for the last five days. Coherent with the experimental data, the starch concentration decreases by 14 — 15%, as seen in Table 4.

**Table 4:** Starch accumulation at 6*h* and 12*h* photoperiods. With a light intensity equal to 160*μmol*· *m*^-2^ · *s*^-1^, we simulate the growth of the plant for 30 days. Then, we measure the starch accumulation for last five days. As noticed in [44], we register a starch decreasing of 14 – 15% as well.

|  |  |  |  |  |  |
| --- | --- | --- | --- | --- | --- |
| Starch concentration at 6h | 48.6601 | 48.5468 | 48.4300 | 48.3097 | 48.1862 |
| Starch concentration at 12h | 57.3064 | 57.0963 | 56.8811 | 56.6614 | 56.4379 |
| Difference (%) | 15.09 | 14.97 | 14.86 | 14.74 | 14.62 |

In [44], the authors find that as photoperiod increases from 6*h* to 12*h*, the starch accumulation significantly decreases. In particular, with a light intensity of 160*μmol* · *m*^-2^ · *s*^-1^, the rate of the accumulation goes from 49.4 ± 2.2*μmolC* · *gFW*^-1^ · *h*^-1^ at 6*h* to 30.9 ± 0.8*μmolC* · *gFW*^-1^ · *h*^-1^ at 12h of light. We simulate the plant growth for 35 days and compute the rate of starch synthesis *r*(*t*) as:

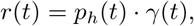

where *p_h_*(*t*) is the hourly production of sucrose by photosynthesis and *γ*(*t*) represents the starch accumulation signal (i.e., the percentage of photosynthesis products devoted to starch production). In Table 5, we collect our results, which are also in this case in agreement with the data provided by [44].

**Table 5:**
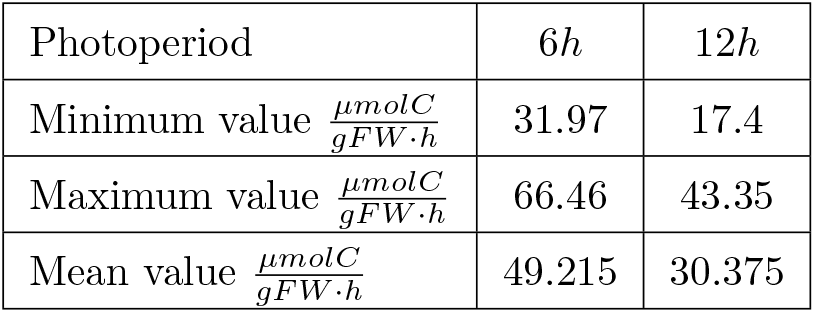
Hourly rate of starch accumulation at 6*h* and 12*h* photoperiods.

| Photoperiod | $6h$ | $12h$ |
| --- | --- | --- |
| Minimum value $\frac{\mu mol C}{gFW \cdot h}$ | 31.97 | 17.4 |
| Maximum value $\frac{\mu mol C}{gFW \cdot h}$ | 66.46 | 43.35 |
| Mean value $\frac{\mu mol C}{gFW \cdot h}$ | 49.215 | 30.375 |

Our model can analyze how plants react to a sudden change in day length. As pointed out by the experiment in [45], plants need time to modify their circadian cycle, and this phenomenon is particularly evident investigating the production of sucrose and starch. In [45], the authors find that passing from short to long days means an increment of starch accumulation (44.4 — 75% higher) and the starch degradation rate (15.44 — 173.4%). Vice versa, passing from long to short days produces the opposite effect, and they measure a significant decrement of starch accumulation (37.5 — 57.14% lower) and the starch degradation rate (62.6 — 73.3%). To reproduce this experiment, we simulate the plant growth for 6 days with 8h of light, and we change the photoperiod to 16h for the following 4 days. Then, we adjust the photoperiod from 16h to 8 for the second comparison. First of all, we notice that - as expected - the plant does not react immediately to the modification of the photoperiod. We show this phenomenon in Figure 2, where we plot the starch dynamics over time. Then, as in [45], we find that increasing the photoperiod generates an increment in starch activity, which decreases otherwise. We collect all the simulation results in Table 6.

**Figure 2:**
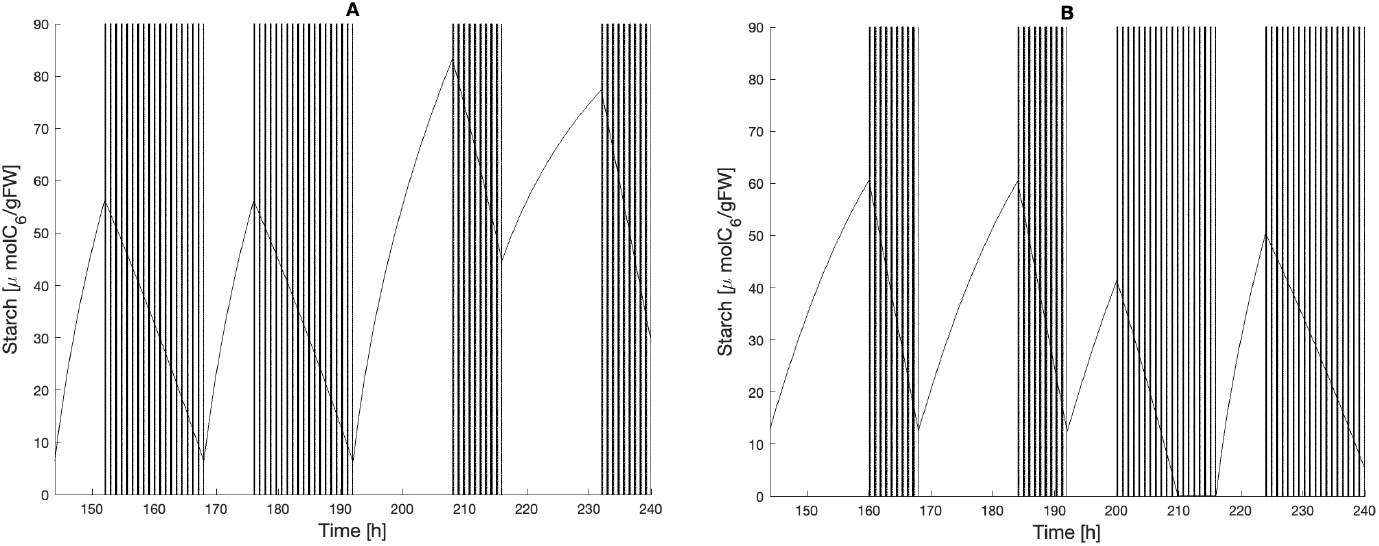
Adaptation of starch in changing photoperiod. We show the starch dynamics 2 days before and 2 days after the photoperiod changes. (A) We plot starch dynamics when passing from short to long days. (B) We plot starch dynamics when passing from long days to short ones.

**Table 6:** Starch dynamics when passing from short days (SD) to long days (LD) and vice versa.

| from SD to LD |  | 8h | 16h | % |
| --- | --- | --- | --- | --- |
| | Starch at dusk $\frac{\mu\text{mol}C_6}{gFW}$ | 55.31 | 81.71 | 47.7 |
| | Starch at dawn $\frac{\mu\text{mol}C_6}{gFW}$ | 6.06 | 43.82 | |
| | Max starch degrad. rate $\frac{\mu\text{mol}C_6}{gFW \cdot h^{-1}}$ | 3.45 | 5.1 | |
| | Min starch degrad. rate $\frac{\mu\text{mol}C_6}{gFW \cdot h^{-1}}$ | 2.33 | 4.29 | |
| | Mean starch degrad. rate $\frac{\mu\text{mol}C_6}{gFW \cdot h^{-1}}$ | 2.89 | 4.69 | 62.3 |
| from LD to SD |  | 16h | 8h | % |
| | Starch at dusk $\frac{\mu\text{mol}C_6}{gFW}$ | 56.06 | 38.2 | -31.85 |
| | Starch at dawn $\frac{\mu\text{mol}C_6}{gFW}$ | 4.79 | 3.38 | -29.4 |
| | Max starch degrad. rate $\frac{\mu\text{mol}C_6}{gFW \cdot h^{-1}}$ | 6.98 | 2.37 | |
| | Min starch degrad. rate $\frac{\mu\text{mol}C_6}{gFW \cdot h^{-1}}$ | 5.9 | 1.53 | |
| | Mean starch degrad. rate $\frac{\mu\text{mol}C_6}{gFW \cdot h^{-1}}$ | 6.44 | 1.95 | -69.7 |

### 5.2. Genetics

We can exploit our model to improve our knowledge of internal plant processes and, in particular, to estimate the effects of genes on specific phenomena.

For instance, in [46], the authors study a mutation in the *Atss3-1* gene of Arabidopsis, which produces an increment amounting to [18.2%, 26.9%] in enzyme activity involved in starch dynamics, with a consequent increment of starch accumulation at dawn (2.5 — 12%) and at dusk (2.6 — 23.2%).

To simulate this behavior, first we need to modify the parameters that affect the starch synthesis, which without mutations are *λ_sdr_* = 0.25 [1/h] and *λ_sdi_* = 0.1 [1/*h*]. To modify these parameters, we simulate 30 days of growth (to avoid dependencies on initial data) at 12h photoperiod in a no-limiting soil condition. Then, we find by trials that the minimum amount of synthesized starch is in the range [18.2%, 26.9%], as outlined by the work [46], when *λ_sdr_* = 0.03 [1/*h*] and *λ_sdi_* = 0.27 [1/*h*]. After setting the new values, we find that our simulation results agree with the data: the starch accumulation at dawn increases by 8.69% and dusk by 12.3%.

### 5.3. Nutrients dynamics

It is well-known that, at fixed soil conditions, plants keep the ratio between nutrients, namely the stoichiometry ratio, almost constant, as also pointed out in [47], and adopt strategies to recover the equilibrium among nutrients [19]. In our model, we consider the plant’s stoichiometry ratio, the correlation with the soil conditions, and the effects of nutrient availability on plant growth.

In particular, using our model, we can estimate the number of days the plant needs to recover the optimal stoichiometry ratio, as we show in Figure 3. Here, we plot the nitrogen-phosphorus ratios (N:P ratios) assuming different initial plant contents in fixed no-limiting soil conditions and considering 12*h* of photoperiod for 30 days of growth.

**Figure 3:**
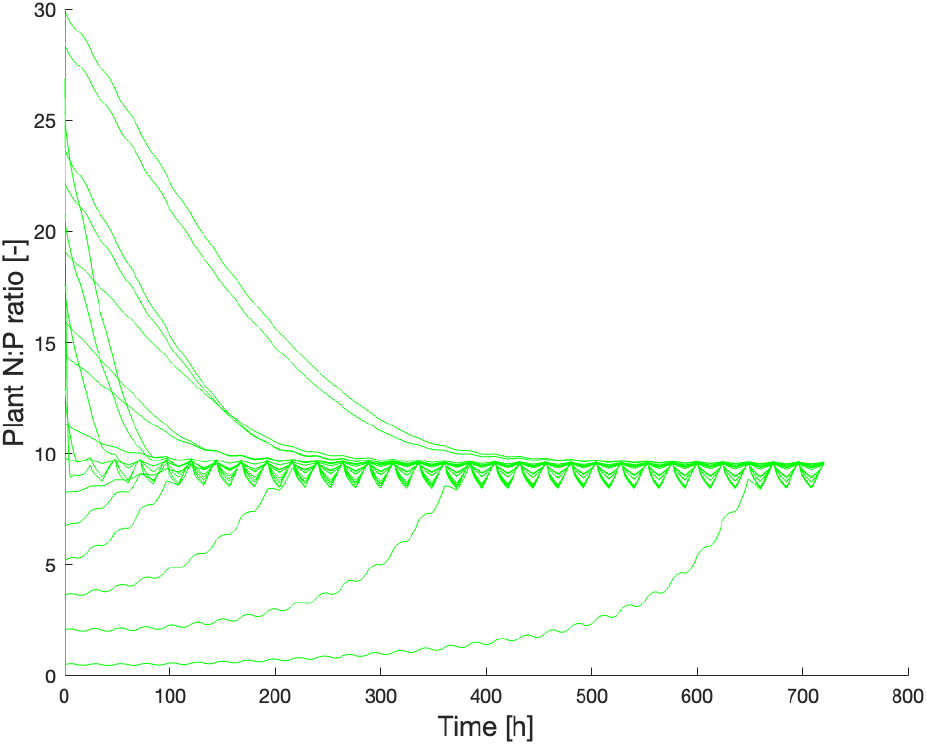
Evolution of N:P ratio from different initial conditions.

The stoichiometry ratio is crucial in agriculture to investigate the concentration of nutrients in the yield after a given cultivation period or to manage fertilizer usage. In [47], they examine the correlation between nutrients contained in soil and the plant and suggest that soil conditions can affect the optimal value of the N:P ratio, as we assume in our model. to verify the accuracy of our results, we test our model considering the experimental data provided by [48]. In this work, the authors grew plants for 58-73 days in 16h photoperiod and tested different soil conditions. To compare their results in terms of growth, they compute the relative growth rate (RGR) as:

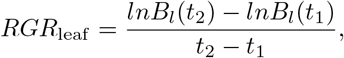

where *t*_2_ is the harvesting time (*t*_2_ ∈ [58,73] days), *t*_1_ is ∈ [42,55] days, and *B_l_* is the leaf biomass. In general, they observe that when nitrogen is the limiting nutrient, an increment of N:P ratios corresponds to an increment in the RGR; when phosphorus is the limiting nutrient, an increment of N:P ratios produces a decrement in the RGR.

To reproduce these experimental observations, we simulate 65 days of growth at 16h photoperiod and compute the RGR at the 48th day. Then, we consider different combinations of concentrations of nitrogen and phosphorus, which we plot in Figure 4. As for the experiments, we obtain the highest values of RGR (top right corner in Figure 4) in a context of limited nitrogen. Moreover, we find the lowest RGR values when the phosphorus concentration is lower, and we notice this behavior observing the plant RGR decreases (moving from top to bottom in Figure 4B) and the N:P ratio increases (moving from top to bottom in Figure 4A).

**Figure 4:**
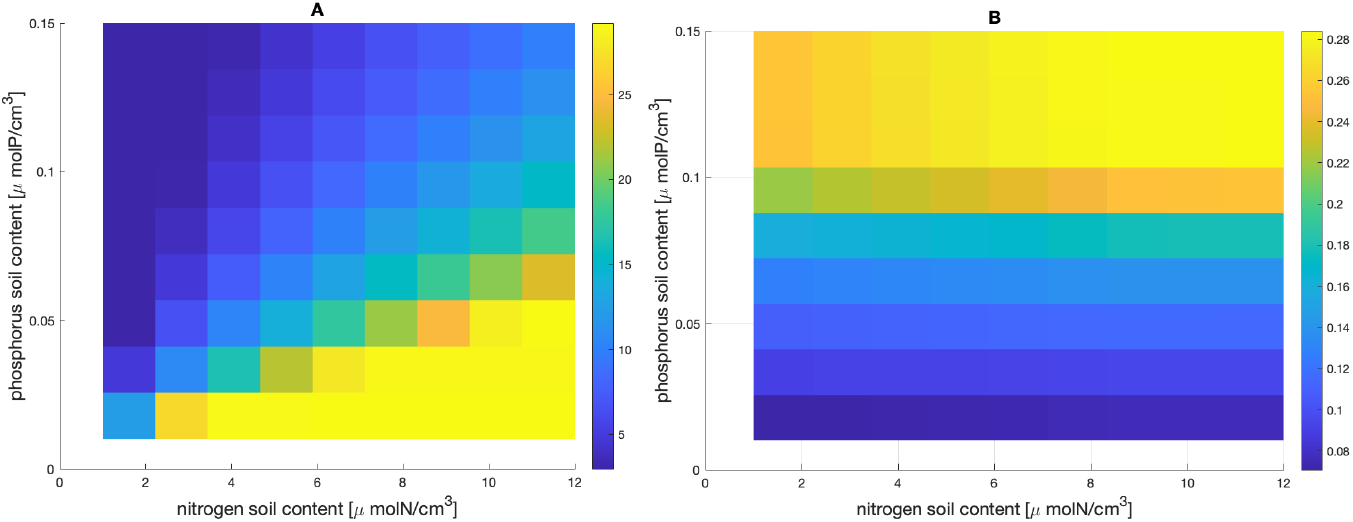
Adaptation of optimal N:P ratio when different soil conditions are experienced. (A) Simulated plant N:P ratio. (B) Simulated leaf RGR fixing *t*_1_ = 48 and *t*_2_ = 65 days.

Considering these results, we think that this model could be profitable in agricultural and ecological studies to estimate how soil characteristics affect the plant’s internal status, improve fertilizer management, and support ecological studies.

### 5.4. The uptake process

As described in Section 2.2.3, we assume that the nutrient uptake rates follow the Michaelis-Menten kinetics, whose parameters are not soil-dependent in our model. Instead, in the work by Narang et al. [21], the authors analyze how the soil state can actually alter these rates, studying two situations: low soil phosphorus (LP, *p*_soil_ = 0.0025*μmolP/cm*^3^), and high soil phosphorus (HP, *p*_soil_ = 0.5*μmolP/cm*^3^). In order to validate our approach, we compare our simulations with their results in Table 7.

**Table 7:**
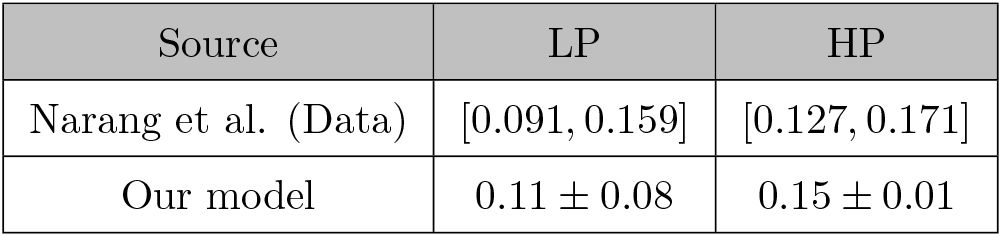
Hourly uptake of phosphorus (P) in poor and rich soil.

As we can notice, this juxtaposition verifies that the affinity signal that we use in our model approximates the uptake rates in a good agreement with the experimental results provided by [21].

### 5.5. Biomass estimation

Our model is a precise tool to estimate biomass production considering soil conditions. In this work done by Trull et al. [49], the authors report two experiments characterized by two soil phosphorus (P) treatments: limited P medium (*p*_soil_^*L*^ = 0.006*μmolP/cm*^3^) and saturated P medium (*p*_soil_^H^ = 1*μmolP/cm*^3^). They grow plants in 16*h* photoperiod for 19 days and measure the total dry biomass (DW) and the root:shoot ratio every 4 days. We use data from the 7th day as the initial condition of our model and compare both fresh biomass and root fraction at the 11th, 15th, and 19th days. To compare fresh and dried biomass, we use the estimation provided in [11], where *FW* = 12.5*DW*. In Table 8, we compare our simulations with the observations provided by [49].

**Table 8:** Comparison of total biomass and root:shoot biomass ratio. We compare our simulations results to the data provided by Trull et al. [49].

| Source | Parameter | Treatment | 11th day | 15th day | 19th day |
| --- | --- | --- | --- | --- | --- |
| Trull et al. (Data) | Root fraction (%) | $p_{\text{soil}}^H$ | 22.2 | 23.1 | 24.7 |
| Our model | Root fraction (%) | $p_{\text{soil}}^H$ | 23.17 | 24.37 | 24.8 |
| Trull et al. (Data) | Root fraction (%) | $p_{\text{soil}}^L$ | 49.4 | 50 | 54.6 |
| Our model | Root fraction (%) | $p_{\text{soil}}^L$ | 43.7 | 50.67 | 54.91 |
| Trull et al. (Data) | Total biomass ($gFW$) | $p_{\text{soil}}^H$ | 0.01675 | 0.0398 | 0.057 |
| Our model | Total biomass ($gFW$) | $p_{\text{soil}}^H$ | 0.0147 | 0.0315 | 0.066 |
| Trull et al. (Data) | Total biomass ($gFW$) | $p_{\text{soil}}^L$ | 0.00287 | 0.00425 | 0.004875 |
| Our model | Total biomass ($gFW$) | $p_{\text{soil}}^L$ | 0.0023 | 0.0038 | 0.0042 |

## 6. Discussion and conclusion

To achieve a global understanding of plant growth, we build a model characterized by the integration of many internal and external biological processes. In particular, we describe the model for *Arabidopsis thaliana*, which we consider as divided in two compartments (i.e., the shoot and the root apparatus of the plant). Then, we simulate its growth considering several factors, such as water availability, light, nitrogen and phosphorus concentrations. In addition, we include starch and sucrose dynamics, photosynthesis, and nutrient uptake, characterizing all these physiological mechanisms as faithfully as possible. We simplify the modelization of the nutrient uptake, designing an internal signals representing the affinity between the plant and the specific nutrient. And we propose a new formalism describing the plant resource allocation, which allows us to test the model on several real scenarios.

This approach, of course, can be considered complex and computationally expensive. On the other hand, we largely show that it guarantees high precision and accuracy in the reproduction of experimental observations. Considering all the main physiological mechanisms involved in plant development in a realistic environment assures us to deal with a more complete and versatile model, which can be applied in different contexts, such as agriculture or genetics.

We show that by applying our model we can investigate how changes in photoperiod and irradiance affect starch and sucrose dynamics. Moreover, we can also analyze how plants react to a sudden change in day length, deeply understanding the starch activity and its accumulation. Remarkably, we can estimate the effects of genes on specific phenomena. Finally, we can simulate different combinations of concentrations of nitrogen and phosphorus to find the optimal condition for the plant. This result is particularly relevant for agriculture and ecology because by understanding how soil conditions affects the plant’s internal status, we can improve fertilizer management and prevent climate change.

As future work, we plan to investigate plant wellness in relation to a specific ecosystem and quantify its efficiency. As shown in [50], it is difficult to have a clear measurement for this feature, for which usually is taken into account only the biomass factor. Thus, we want to use our comprehensive model to formalize this notion.

## Acknowledgment

This project has received funding from the European Union’s Horizon 2020. Research and Innovation Programme under Grant Agreement No 824074.

