## Supplementary Material for "A mechanistic model towards ecological inclusion for a better plant growth understanding"

Lucia Nasti*

*Gran Sasso Science Institure, L'Aquila, Italy*

Emanuela Del Dottore*

*Bioinspired Soft Robotics laboratory, Istituto Italiano di Tecnologia, Genova, Italy*

Fabio Tedone

*Gran Sasso Science Institute, L'Aquila, Italy*

Michele Palladino

*Gran Sasso Science Institute, L'Aquila, Italy*

*Università dell'Aquila, L'Aquila, Italy*

Barbara Mazzolai

*Bioinspired Soft Robotics laboratory, Istituto Italiano di Tecnologia, Genova, Italy*

Pierangelo Marcati

*Gran Sasso Science Institute, L'Aquila, Italy*

---

### 1. Model of plant growth

The model consists of 10 non-linear ordinary differential equations, which we are going to define in detail. All the parameters are summarized in Section. We assume that the biomass is measured in grams of fresh weight ($[gFW]$), while the sugar (sucrose or starch) is measured in $\mu molC_6$. In some equations, we introduce a small correction factor ($\epsilon = 1{-}20$), which prevents discontinuities

*Corresponding authors

in the model when some variable go to 0. It is a numerical adjustment that does not affect the results of the simulations.

#### *1.1. Photosynthesis*

We assume the plant is composed of roots and leaves, so that the photosynthetically active biomass is the leaf biomass $b_l$ $[gFW]$. We assume that:

$$p_h(t) \propto \frac{b_l}{b_l + \theta_{bl}}. \tag{1}$$

Photosynthesis depends on water availability. Named $w_{soil}(t)$ $[-]$ the volumetric water content at time $t$ and $\theta_w$ $[-]$ a parameter to be estimated, the water stress is modelled by the saturating function:

$$\bar{w}(t) = \frac{w_{soil}(t)}{w_{soil}(t) + \theta_w} \; [-]. \tag{2}$$

$$\bar{w}(t) = \frac{w_{soil}(t)}{w_{soil}(t) + \theta_w}. \tag{3}$$

The rate of carbon production depends on plant's nitrogen $n(t)$ and phosphorus $p(t)$. According to [1], we assume that the relation between the carbon and nitrogen is the following:

$$15.44 \frac{gC}{gN \cdot N} = 2.21 \frac{\mu mol C_6}{\mu mol N \cdot h}.$$

Hence, the minimum quantity of nitrogen required to sustain the maximum rate of photosynthesis is:

$$n_{\text{ph}} = \frac{p_h^{\max}}{2.21} \tag{4}$$

Therefore, we assume that:

$$p_h(t) \propto \bar{n}(t) = \max(0, \frac{2n(t)}{n(t) + n_{\text{ph}}} - 1). \tag{5}$$

When $n(t) < n_{\text{ph}}$), $\bar{n}(t) = 0$, thus the photosynthesis cannot start because there is not enough nitrogen. Otherwise, when $n(t) >> n_{\text{ph}}$, $\bar{n}(t) \approx 1$, hence the photosynthesis is not limited by the nitrogen concentration.

Similarly, we can model the effects of phosphorus on photosynthesis as follows:

$$p_h(t) \propto \bar{p}(t) = \max(0, \frac{2p(t)}{p(t) + p_{\text{ph}}} - 1). \tag{6}$$

We assume that:

$$p_{\text{ph}} = \frac{n_{\text{ph}}}{\bar{O}},$$

where $\bar{O}$ represents the ratio between nitrogen and phosphorus in a no-limiting soil conditions (estimated from [2]). According to the law of minimum, the most limiting resource affects the photosynthesis, namely

$$p_h(t) \propto min(\bar{n}(t), \bar{p}(t)). \tag{7}$$

In addition, we model the situation in which the concentration of phosphorus in soil is toxic for the plant, causing a reduction of photosynthesis of 50% [3]:

$$p_h(t) \propto p_{\text{tox}}(t, p_{\text{soil}}(t)) = \delta_{pt} + (1 - \delta_{pt}) \min\left(1, \mu_1 \exp(\mu_2 p_{\text{soil}})\right), \tag{8}$$

where $\delta_{pt} = 0.5$ represents the maximum reduction possible, and $\mu_1$ and $\mu_2$ are parameters that have to be estimated.

We include the *photosynthetic control function* $C(t, a(t))$ to model the interaction between the photosynthesis rate and the starch accumulation. Indeed, when the starch accumulated is enough to sustain the plant metabolism, the photosynthesis rate can be reduced:

$$p_h \propto C(t, a(t)) = \lambda_c + (1 - \lambda_c)\frac{a^{\max}(t) - a(t)}{a^{max}(t)}, \tag{9}$$

where $\lambda_c$ is a parameter to estimate and ensures that a minimum amount of photosynthesis is still produced to sustain the daily respiration. The function $a(t)$ is the plant starch content and $a^{\max}$ is the maximum amount of starch that the plant needs during the night. The parameter $\tau^{\max}$ represents the maximum nightly degradation of starch, which can be estimated by literature. Then:

$$a^{\max} = \tau^{\max}(24 - f), \tag{10}$$

where $f$ is the photoperiod.

We include the influence of water ($w_{\text{soil}}(t)$) using the following saturation function:

$$\bar{w}(t) = \frac{w_{\text{soil}}(t)}{w_{\text{soil}}(t) + \theta_w}.$$

.

To summarise, the photosynthesis directly depends on the plant's starch content $a(t)$, the nitrogen content $n(t)$, the phosphorus content $p(t)$, the water availability $w_{\text{soil}}(t)$, the possibly toxicity of soil phosphorus content $p_{\text{soil}}(t)$ and the leaf biomass $b_l(t)$:

$$p_h(t) = p_h^{\max} L(t)\bar{w}(t)\min(\bar{n}(t), \bar{p}(t))p_{\text{tox}}(t, p_{\text{soil}}(t))C(t, a(t))\frac{b_l(t)}{b_l(t) + \theta_{bl}}. \tag{11}$$

*1.2. Starch and sucrose*

The photosynthetic products are accumulated as starch ($a(t)$) to sustain the nocturnal metabolism, and as sucrose ($s(t)$) to sustain the diurnal metabolism. We introduce the *starch accumulation signal* $\gamma(t)$ to model the starch synthesis:

$$\frac{d\gamma(t)}{dt} = L(t)\left(-\gamma(t)\lambda_{\text{sdr}}\frac{s^{\min}}{s(t) + s^{\min}} + (1 - \gamma(t))\lambda_{\text{sdi}}\frac{s(t)}{s(t) + s^{\max}}\right) + (1 - L(t))(1 - \gamma(t))\lambda_{sni}\frac{s^{\min}}{s^{\min} + s(t)}, \tag{12}$$

where $\lambda_{\text{sdr}}$, $\lambda_{\text{sdi}}$, $\lambda_{\text{sni}}$ are parameters to estimate. From literature, we estimate an upper bound ($s^{\max}$) and a lower bound ($s^{\min}$) for the sucrose content $s(t)$. In case $s(t) > s^{\max}$, the plant is accumulating sucrose, then there is an inhibition of sucrose synthesis and a starch accumulation, which we model using a saturation function:

$$(1 - \gamma(t))\lambda_{\text{sdi}}\frac{s(t)}{s(t) + s^{\max}}.$$

As a consequence, the function $\gamma(t)$ increases because of the daily sucrose accumulation. In case $s(t) < s^{\min}$ during the day, the plant is daily starving because of fast sucrose consumption that triggers sucrose synthesis and inhibits starch accumulation [4]. This phenomenon is modelled by the following saturation function:

$$-\gamma(t)\lambda_{\text{sdr}}\frac{s^{\min}}{s(t)+s^{\min}},$$

which reduces the function $\gamma(t)$. Instead, when $s(t) < s^{\min}$ during the night, it means that the starvation is experienced because not enough starch has been accumulated. Then, the next day the plant will need to increase the amount of stored starch [5]. Therefore, the saturation function

$$(1-\gamma(t))\lambda_{\text{sni}}\frac{s^{\min}}{s^{\min}+s(t)}$$

increases $\gamma(t)$ and, hence, the starch will be synthesized the next day.

Given $\gamma(t)$, we can estimate the starch and sucrose dynamics. The following equation describes the starch dynamics:

$$\frac{da(t)}{dt} = \gamma(t)p_h(t) - \tau_{\text{as}}(t), \tag{13}$$

where $\gamma(t)p_h(t)$ estimates the daily rate of starch synthesis, $\tau_{\text{as}}$ describes the nightly rate of starch degradation and conversion in sucrose. In general, plants convert in sucrose almost all the starch accumulated during the day, except for a minimum quantity ($a^{\min}$), which can be estimated by literature. In addition, the starch degradation should be in almost linear manner, meaning that it should take into account the time remaining before the next dawn. We assume the following formulation:

$$\tau_{as}(t) = \begin{cases} \min\left(\tau^{\max}, (1-L(t))\frac{a^{\text{dusk}}(t)-a^{\min}}{24-f}\left(\frac{t}{24}+\left(1-\frac{t}{24}\right)\left(1-\frac{s(t)}{s(t)+s^{\max}}\right)\right)\right) & a(t) \geq a^{\min} \\ 0 \qquad a(t) < a^{\min} \end{cases}, \tag{14}$$

where $a^{\mathrm{dusk}}(t) = a(f^{\mathrm{real}}$ is the plant's starch content at the end of the light period (namely when $t = f^{\mathrm{real}}$, being $f^{\mathrm{real}}$ the real photoperiod. The term:

$$(1 - L(t))\frac{a^{dusk}(t) - a^{min}}{24 - f(t)}$$

means that starting from $a^{\mathrm{dusk}}$ the starch is consumed up to $a^{\mathrm{min}}$ during the night period. We notice that the quantity of starch consumption depends on the *real* photoperiod (since the starch degradation starts as soon as the dark period starts), while the rate of starch degradation depends on the *perceived* photoperiod $f$, since the starch degradation depends on the plant's circadian clock. The term:

$$(\frac{t}{24} + (1 - \frac{t}{24})(1 - \frac{s(t)}{s(t) + s^{\mathrm{max}}}))$$

reduces the starch degradation rate if the sucrose $s(t)$ is accumulating. This reduction depends on the time before the expected dawn. When $\frac{t}{24}$ is smaller, the night period is still long, then the sucrose accumulation affects $\tau_{\mathrm{as}}$. When $t \approx 24$, sucrose accumulation does not affect the starch degradation since the day is starting and the sucrose would be necessary for the plant metabolism. We consider $\tau^{\mathrm{max}}$ as the maximum rate for the starch degradation.

As stated in the main article, the sucrose dynamics is complex since it considers maintenance and growth respiration that we formulate respectively as follows:

$$r^{\mathrm{m}}(t) = \bar{r}_m(t)s(t) \tag{15}$$

$$r^{\mathrm{g}}(t) = \bar{r}_g(t)s(t), \tag{16}$$

both proportional to sucrose content.

The maintenance proportional function $\bar{r}_{\mathrm{m}}(t)$ is defined as follows:

$$\bar{r}_m(t) = r_m^0\chi_{\mathrm{np}}(t) + \theta_s \ , \tag{17}$$

which can be divided into two terms:

- $r_m^0$ is a frequency term for the active loading of sucrose. while $\chi_{\rm np}(t)$ represents the influence of nutrients concentration on plants growth;
- $\theta_s$ that is a frequency parameter, simulating the losses of sucrose during the transport because of the porous structure of the phloem.

Therefore, we have $\bar{r}_m(t) = \bar{r}_m(t)(t, r_m^0, \theta_s)$.

According to [5], when the plant experiences nocturnal starvation, the growth respiration is reduced the next day. In our model, the nightly starvation increases the daily starch signal $\gamma(t)$ and, consequently, the growth respiration function $\bar{r}_g(t)$ is reduced when $\gamma(t)$ increases. Moreover, as shown in [6], there are many phenomena that can reduce the growth stimulus. Indeed, we can write $\bar{r}_g(t) = \bar{r}_g(t, \gamma(t), \eta(t))$, where $\eta(t)$ is defined as follows:

$$\eta(t) = (\delta_{\rm npd} + (1 - \delta_{\rm npd})\bar{\eta}_{\rm def}(t))\, \bar{\eta}_{\rm tox}(t).$$

In addition we include the function $\eta_{\rm def}(t)$, to model the influence of nutrients on growth, and the function $\eta_{\rm tox}(t)$ to model the excess of nitrogen causing growth reduction. Therefore, we have:

$$\bar{\eta}_{\rm def}(t) = \min\left(1, \mu_3 \exp\left(\mu_4 n_{\rm soil}\right)\right) \min\left(1, \mu_5 \exp\left(\mu_6 p_{\rm soil}\right)\right); \tag{18}$$

$$\bar{\eta}_{\rm tox}(t) = \delta_{\rm nt} + (1 - \delta_{\rm nt}) \min\left(1, \mu_7 \exp(\mu_8 n_{\rm soil})\right). \tag{19}$$

$$\tag{20}$$

Here, $\delta_{\rm npd} = 0.3$ and $\delta_{\rm nt} = 0.7$ are taken from [6]. The other parameters ($\mu_3$, $\mu_4$, $\mu_5$, $\mu_6$) have to be estimated.

Besides sucrose, respiration consumes both nitrogen and phosphorus [7], hence the respiration should be limited if there are not enough nutrients. In [1], the grams of nitrogen consumed for each gram of carbon used during the respiration is equivalent to

$$c_{\rm sn} = \frac{1}{4.06} \frac{gN}{gC} = 1.724 \ \left[\frac{\mu mol N}{\mu mol C_6}\right].$$

And, for the phosphorus, we assume:

$$c_{\mathrm{sp}} = \frac{c_{\mathrm{sn}}}{\bar{\mathcal{O}}} \left[\frac{\mu mol P}{\mu mol C_6}\right]. \tag{21}$$

If the plant's nutrient contents are not enough to sustain the respiration, then the metabolism is reduced according to the following equation:

$$\chi_{\mathrm{np}}(t) = min\left(1, \frac{n(t)}{s^{\max} c_{\mathrm{sn}}}\right) min\left(1, \frac{p(t)}{s^{\max} c_{\mathrm{sp}}}\right).$$

Then, the function of growth respiration is the following:

$$\bar{r}_g(t) = r_g^0\big(\lambda_g + (1-\lambda_g)\,(1 - L(t)\gamma(t))\big)\eta(t)\chi_{\mathrm{np}}(t) \tag{22}$$

where $r_g^0$ is the frequency parameter which simulates the rate of sucrose devoted to produce new biomass; $\lambda_g$ measures the strength of the nocturnal starvation in reducing the growth respiration. Since sucrose is consumed also by carbon transportation along the plant, we add the following function:

$$r^t(t) = \bar{r}_t\left(r^m(t) + r^g(t)\right).$$

In [8], the authors estimate $\bar{r}^u$, namely the grams of carbon necessary to uptake one gram of phosphorus. According to their results, we can estimate the sugar cost of uptake phosphorus as:

$$p_c = 0.053\frac{\mu g Sugar}{\mu g P} \approx 0.053\frac{\mu g Sucrose}{\mu g P} = 0.585\frac{\mu mol C_{12}H_{22}O_{11}}{\mu mol P} \approx 0.29\frac{\mu mol C_6}{\mu mol P}. \tag{23}$$

And, since there is no reason to assume a different cost for nitrogen, we assume the sugar cost of uptake nitrogen is:

$$n_c = 0.053\frac{\mu g Sugar}{\mu g N} \approx 0.053\frac{\mu g Sucrose}{\mu g N} = 1.296\frac{\mu mol C_{12}H_{22}O_{11}}{\mu mol N} \approx 0.65\frac{\mu mol C_6}{\mu mol N}. \tag{24}$$

Named $u_n(t)$, the rate of nitrogen uptaken at time $t$ and $u_p(t)$ the rate of phosphorus uptaken at time $t$, the rate of sucrose consumption due to the uptake will be:

$$r^u(t) = n_c u_n(t) + p_c u_p(t).$$

Finally, all the processes sucrose consuming can be collected in the following sucrose dynamics:

$$\frac{ds(t)}{dt} = (1-\gamma(t))\, p_h(t) + \tau_{as}(t) - r^u(t) - r^t(t) - r^m(t) - r^g(t). \tag{25}$$

*1.3. Sucrose allocation and growth*

As mentioned in the main article, plants are able to divide the sucrose devoted to growth between leaves and roots. We use $f_r(t) \in [0,1]$ to define the portion of $r^g(t)$ consumed by the roots, while $1 - f_r(t)$ is the portion of sucrose allocated to the leaf biomass. The amount of sucrose that the plants invest in root biomass is influenced by the *root's activity*, which depends on the nutrients availability and sucrose content [9, 10, 11].

In particular, the root's priority increases when any nutrient is limiting, for instance when plant's nutrient content is not sufficient to sustain the plant's metabolism. Instead, the roots' activity decreases in rich soils where the nutrients are easily to uptake. Concerning the uptake of phosphorus, it seems that the root's activity increases when the soil phosphorus content increases and becomes toxic. In this case, the plant uptakes more phosphorus than necessary: this phenomenon is known as *luxury uptake* of phosphorus [12] and, in our case, it is modelled by the function $p_{\text{lux}}(t, p_{\text{soil}})$. This function depends on the phosphorus content $p_{\text{soil}}$, which is estimated by [13].

The root's activity decreases if the plant's sucrose content or the nutrients concentrations are limiting. Therefore the fuction describing the root's activity is the following:

$$\frac{df_r(t)}{dt} = p_{\text{lux}}(t)(1-f_r(t))\left(a_n(t)f_n(t) + (1-f_n(t))a_p(t)\right) - \tag{26}$$

$$- f_r(t)\left(\frac{n(t)(1-f_n(t))}{n(t)+n^{\min}(t)} + \frac{p(t)f_n(t)}{p(t)+p^{\min}(t)} + \frac{s^{\min}}{s(t)+s^{\min}}\right). \tag{27}$$

The first term of the function above

$$p_{\text{lux}}(t)(1-f_r(t))\left(a_n(t)f_n(t) + (1-f_n(t))a_p(t)\right)$$

aims to increase the root's priority $f_r(t)$ up to its maximum value 1. The increment is driven by the nitrogen and phosphorus uptake activities $a_n(t) \in [0,1]$ and $a_p(t) \in [0,1]$, respectively. Let us note that $a_n(t),\ a_p(t)$ are not the uptake rates of nutrients from the soil but two signals that regulate the plant's affinity to each nutrient. We recall that $a_n(t),\ a_p(t)$ increase when the nutrient (nitrogen or phosphorus) is limiting from the plant's perspective, namely when the plant's nutrient content does not sustain the actual metabolism. Therefore, the increment of $f_r(t)$ is proportional to $a_n(t)$ and $a_p(t)$, through the proportional function $f_n(t) \in [0,1]$. The function $f_n(t)$ estimates the role of the stoichiometry N:P ratio on the root's priority. Let $\mathcal{O}$ the N:P ratio that plants tend to keep constant in given soil conditions. According to [14], $\mathcal{O}$ is proportional to the soil N:P ratio. We assume that

$$\mathcal{O}(t) = \min\left(\max\left(\lambda_O \bar{\mathcal{O}} \frac{n_{\text{soil}}(t)}{p_{\text{soil}}(t)}, \mathcal{O}^{\min}\right), \mathcal{O}^{\max}\right), \tag{28}$$

where $n_{\text{soil}}$ and $p_{\text{soil}}(t)$ are the soil nitrogen and phosphorus contents at time $t$, $\lambda_O$ is a parameter to be estimated, $\bar{\mathcal{O}}$ is the plant N:P ratio in no limiting soil conditions [2] and $\mathcal{O}^{\min},\ \mathcal{O}^{\max}$ are the minimum and maximum values for $\mathcal{O}$ (estimated by [14]). Therefore, we define

$$f_n(t) = \frac{\mathcal{O}(t)}{\mathcal{O}(t) + \frac{n(t)}{p(t)+\varepsilon}} \in [0,1] \tag{29}$$

When the plant N:P ratio is close to the value that the plant tries to keep constant in the given soil conditions (namely $n(t)/p(t) \approx \mathcal{O}(t)$), $f_n(t) \approx 0.5$ and the nutrients have the same weight in defining the root's priority $f_r(t)$. When the stoichiometry ratio is not close to the value expected by the plant ($f_n \to 1$ if $n(t)/p(t) < \mathcal{O}(t)$ or $f_n \to 0$ if $n(t)/p(t) > \mathcal{O}(t)$), then the weight of the limiting nutrient increases in defining the root's priority. For example, assume $n(t)/p(t) << \mathcal{O}(t)$. Therefore $f_n(t) \approx 1$. Any activity of the roots to uptake nitrogen ($a_n(t)$) greatly affects the root's priority $f_r(t)$. Any activity of the roots to uptake phosphorus ($a_p(t)$) has smaller effects on the root's priority ($1 - f_n(t) \approx 0$) since the phosphorus is not as limiting as the nitrogen.

The second right-hand side term

$$-f_r(t)\left(\frac{n(t)(1-f_n(t))}{n(t)+n^{\min}(t)}+\frac{p(t)f_n(t)}{p(t)+p^{\min}(t)}+\frac{s^{\min}}{s(t)+s^{\min}}\right)$$

aims to decrease the root's priority when the plant's nutrient contents can sustain the internal metabolism and the sucrose content cannot sustain the respiration. Indeed, the minimum thresholds of nitrogen ($n^{\min}(t)$ ) and phosphorus ($p^{\min}(t)$ ) contents to sustain the actual metabolism. When enough nutrient is stored ($n(t) >> n^{min}(t)$ or $p(t) >> p^{\min}(t)$) the root's priority decreases accordingly. The same holds when the sucrose is lmiting. When the sucrose content is below the starvation threshold $s^{\min}$, the root's priority decreases so that more sucrose is devoted to the leaf growth. Let us note that the stoichiometry ratio can affect the decrement of $f_r(t)$. For example, assume enough nitrogen is stored ($n(t) >> n^{\min}(t)$) and that the stoichiometry ratio is not satisfied ($n(t)/p(t) << \mathcal{O}(t)$ and $f_n(t) \approx 1$). Since the nitrogen is accumulated in the plant ($n(t)/(n(t)+n^{\min}(t)) \approx 1$), it is expected that the root's priority decreases. Nevertheless, this decrement is not experienced. Indeed, since $f_n(t) \approx 1$, the nitrogen is limiting for the plant (meaning that the stoichiometry ratio is not satisfied because of nitrogen lack). Consequently, $1 - f_n(t) \approx 0$ and $n(t)(1-f_n(t))/(n(t)+n^{\min}(t)) \approx 0$ and $f_r(t)$ is not affected by nitrogen accumulation in the plant.

Finally, we model the sucrose conversion in new biomass for the growth. The rate of sucrose devoted to the production of new root biomass will be proportional to $f_r(t)r^g(t)b_l(t)$ , namely the percentage devoted to roots of the total sucrose produced in the total leaf biomass and loaded into the phloem for the growth. Recall that by $b_l(t)$ and $b_r(t)$ we mean the total leaf and root biomass respectively.

In addition, as noted in [15], an excess of nitrogen in soil could induce a reduction in the primary root growth and consequently, a fast development of leaves. Let $n_{\text{tox}}(t) \in [0,1]$ be the function that simulates this phenomenon:

$$n_{\text{tox}}(t) = \min\left(1, \mu_9 \exp(\mu_{10} n_{\text{soil}})\right). \tag{30}$$

$\mu_9$ and $\mu_10$ are parameters to be estimated that measure the intensity of the negative effect of the nitrogen soil content on the root growth. Two phenomena negatively affect both leaves and roots growth: the death of tissues and the competition in limiting environments. Let $\theta_{\text{ld}}$ and $\theta_{\text{rd}}$ be the death rates of leaf and root tissues respectively. As revised in [16], an excess of water into the soil reduces the oxygen available in the above-ground biomass and increases the death of leaf tissues. Therefore, the death rate of leaf tissues will be proportional to a parameter $\theta_{\text{w2}}(t)$ that measures the excess of water into the soil. We assume $\theta_{\text{w2}}(t) = 1$ when there is no excess of water. Finally, let $\theta_{\text{lc}}$ and $\theta_{\text{rc}}$ two parameters that estimate the self-shading in leaves and the overproduction of roots, respectively. Therefore, we can model the growth of tissues as:

$$\frac{db_l}{dt} = \lambda_{\text{sb}}(1 - f_r(t)n_{\text{tox}}(t))r^g(t)b_l(t) - \theta_{\text{w2}}(t)\theta_{\text{ld}}b_l(t) - \theta_{\text{lc}}b_l^2(t), \tag{31}$$

$$\frac{db_r}{dt} = \lambda_{\text{sb}}f_r(t)n_{\text{tox}}(t)r^g(t)b_l(t) - \theta_{\text{rd}}b_r(t) - \theta_{\text{rc}}b_r^2(t), \tag{32}$$

where $\lambda_{\text{sb}}$ is a parameter to be estimated that measures the sucrose conversion in biomass.

#### *1.4. Uptake of nutrients*

The uptake rates of nitrogen and phosphorus can be estimated by using the Michaelis-Menten kinetics:

$$u_{n\text{MM}}(I_n, k_n) = I_n \frac{n_{\text{soil}}}{n_{\text{soil}} + k_n}, \quad u_{p\text{MM}}(I_p, k_p) = I_p \frac{p_{\text{soil}}}{p_{\text{soil}} + k_p}, \tag{33}$$

where $I_n$, $k_n$, $I_p$ , $k_p$ are called Michaelis-Menten parameters and their values are thought to be soil dependent. Instead, we assume these parameters are fixed and not affected by soil conditions.

In particular, the maximum values observed in [17, 18] ($I_n^{\max}$, $k_n^{\max}$, $I_p^{\max}$, $k_p^{\max}$) are used to estimate these fixed values. Then, we assume that the uptake rate $u_{n\text{MM}}$ and $u_{p\text{MM}}$ (when the Michaelis Menten parameters are fixed) could be reduced by the plant according to the plant's internal status. We define an internal signal $a_n(t) \in [0, 1]$ for the nitrogen and an internal signal $a_p(t) \in [0, 1]$ for the phosphorus that simulates the affinity of the plant to that nutrient.

When $a_n(t) = 1$ or $a_p(t) = 1$ the plant has the maximum affinity to nitrogen and phosphorus and the uptake rate is equal to $u_{n\mathrm{MM}}$ and $u_{p\mathrm{MM}}$. When $a_n(t) = 0$ or $a_p(t) = 0$, the uptake rate of that nutrient is turned off even if the nutrient is available into the soil. Finally, since the uptake is a sucrose-consuming process, the hourly uptake rate is reduced if not enough sucrose is available, according to the following saturating functions:

$$u_n^{\mathrm{sat}}(t) = \frac{s(t)}{s(t) + n_c u_{n\mathrm{MM}}(t) a_n(t) \lambda_t + \varepsilon}, \quad u_{\mathrm{sat}}^{p}(t) = \frac{s(t)}{s(t) + p_c u_{p\mathrm{MM}}(t) a_p(t) \lambda_t + \varepsilon}. \tag{34}$$

,

where the paramenter $\lambda_t = 1h$ is fixed since we are considering the hourly uptake rate.

To summarise, the total nitrogen and phosphorus hourly uptaken by the roots will be:

$$u_n(t) = u_{n\mathrm{MM}}(I_n^{\max}, k_n^{\max}) a_n(t) u_n^{\mathrm{sat}}(t) \frac{b_r(t)}{b_l(t) + b_r(t)}, \tag{35}$$

$$u_p(t) = u_{p\mathrm{MM}}(I_p^{\max}, k_p^{\max}) a_p(t) u_p^{\mathrm{sat}}(t) \frac{b_r(t)}{b_l(t) + b_r(t)}. \tag{36}$$

We notice that the uptake rate is still dependent on the soil conditions, but we do not need to estimate the Michaelis-Menten parameters each time the nutrient soil content changes. Therefore, the nutrients are distributed along leaves and roots and consumed by processes like the the respiration and the photosynthesis. Additionally, the starch requires a maintenance respiration (and then a cost in nutrients) due to the production of storage structures [19]. We assume the following total cost for the nitrogen:

$$c_n(t) = \lambda_{\mathrm{f1}} \left( r^m(t) + r^g(t) + \bar{r}_m(t) a(t) \right) + p_h(t) \lambda_{\mathrm{f2}}, \tag{37}$$

being $\lambda_{\mathrm{f1}}$, $\lambda_{\mathrm{f2}}$ two proportionality parameters to be estimated. As usual, we assume

$$c_p(t) = \frac{c_n(t)}{\bar{\mathcal{O}}}. \tag{38}$$

Finally, nitrogen and phosphorus contents will vary according to:

$$\frac{dn(t)}{dt} = u_n(t) - c_n(t), \tag{39}$$

$$\frac{dp(t)}{dt} = u_p(t) - c_p(t). \tag{40}$$

It remains to describe the affinity signals $a_n(t)$ and $a_p(t)$. The main difficult is to define *enough nutrients stored* in a quantitative manner. Then, we estimate four thresholds: two lower bounds $n^{\min}(t)$ and $p^{\min}(t)$ and two upper bounds $n^{\max}(t)$ and $p^{\max}(t)$ for the plant's nitrogen and phosphorus contents, respectively. The lower bounds $n^{min}(t)$ and $p^{min}(t)$ estimate the amount on nitrogen and phosphorus necessary to sustain the metabolism, at least for one day and in the worst conditions. For the nitrogen, we assume

$$n^{\min}(t) = \left(24\lambda_{f1}\left(r_m^0\left(s^{\max} + a^{\max}(t)\right) + r_g^0 s^{\max}\right) + f(t)\lambda_{\mathrm{f2}} p_h^{\max}\right)\frac{1}{\bar{\eta}_{\mathrm{def}}(t)}. \tag{41}$$

Both sucrose and starch consume nitrogen during the maintenance respiration. The worst case for the maintenance and growth respiration is when both $s(t) = s^{\max}$ and $a(t) = a^{\max}$ for the whole day (24 hours). The worst case for the photosynthesis is when $p_h(t) = p_h^{\max}$ for the whole perceived day-length $f(t)$. Finally, the function $\bar{\eta}_{\mathrm{def}}(t)$ simulates the increased stimulus to store more resources in poor soils. For the phosphorus, we assume:

$$p^{\min}(t) = \frac{n^{\min}}{\bar{\mathcal{O}}}. \tag{42}$$

The upper bounds $n^{\max}(t)$ and $p^{\max}(t)$ are computed according to the memory concept proposed in [20]. In this paper, it is proposed that the plant stores nutrients up to a maximum threshold. This threshold depends on the number of days the plant could sustain the actual metabolism if suddenly the soil nutrients are not anymore available. The number of days $D$ $[-]$ is a species-specific number and, for herbaceous plants, it is estimated as $D = 4$ $[-]$.As a consequence:

$$n^{\max}(t) = Dn^{\min}(t); \tag{43}$$

$$p^{\max}(t) = Dp^{\min}(t). \tag{44}$$

We can now define the affinity signals as follows:

$$\frac{da_n(t)}{dt} = (1 - a_n(t)) \left( \left( 1 - \frac{u_n(t)}{u_n(t) + c_n(t) + \varepsilon} \right) \frac{n^{\max}(t)}{n^{\max}(t) + n(t)} + \frac{p_h(t)}{p_h^{\max}} - a_n(t)\lambda_k(1 - 2f_n(t)) \right) - \\ -a_n(t) \left( \frac{n(t)}{n(t) + n^{\min}} + \frac{n_c u_n(t)}{n_c u_n(t) + p_h(t)(1 - \gamma(t)) + \tau_{\text{as}}(t) + \varepsilon} \right); \tag{45}$$

$$\frac{da_p(t)}{dt} = (1 - a_p(t)) \left( \left( 1 - \frac{u_p(t)}{u_p(t) + c_p(t) + \varepsilon} \right) \frac{p^{\max}(t)}{p^{\max}(t) + p(t)} + \frac{p_h(t)}{p_h^{\max}} + a_p(t)\lambda_k(1 - 2f_n(t)) \right) - \\ -a_p(t) \left( \frac{p(t)}{p(t) + p^{\min}} + \frac{p_c u_p(t)}{p_c u_p(t) + p_h(t)(1 - \gamma(t)) + \tau_{\text{as}}(t) + \varepsilon} \right). \tag{46}$$

$\lambda_k$ is a parameter to be estimated that measures the strength of the stoichiometry signal in defining the plant's affinity.

### 2. Parameter estimation

As stated in the main article, we estimate the parameters from the works done in [21, 13, 6].

In [21], the authors grew *Arabidopsis* under 5 different photoperiods ($4h$, $6h$, $8h$, $12h$, $18h$ of light) for 29 days, assuming non-limiting medium growth. From their results we obtained seven fixed values and 3 parameters dependent on the photoperiod, being the plant development affected by the thermal time [22]. To verify the correctness of the fitting, we firstly compared the dynamics of sucrose and starch content during the day. The starch dynamics simulated by the model follows the starch content values of [21] for the most of the time (figures 1A-C and figure 1A-B). The behaviour of the sucrose from numerical simulations can be considered in agreement on average with the biological data, even though less match is found between the peaks at dawn and dusk. In fact, the oscillations in sucrose content are smoothed in the model. For instance, from the $4h$ photoperiod (figure 1D), we can observe a single higher peak in sucrose dynamics during night obtained from our model instead of two as for the biological data. A smoother behaviour is more evident in $18h$ of light (figure

2D); a closer view on the oscillations for this photoperiod is reported in figure 3A. Another defect concerns the delay in the numerical results with respect to the experimental data. For instance in the highest peak in the 18$h$ photoperiods, the delay is approximately of 6$h$ (figure 3A), while in the 8$h$ photoperiods (figure 3B), there is a delay of 4$h$. Such a delay can be due to the starch degradation equation, whose formulation requires further investigations. In fact, both the delay and the smoother dynamic effects could imply an on/off mechanism with respect to some thresholds, that should firstly be investigated from biological experimentation.

Data in [13, 6] are used to estimate effects of toxic and limit soil nutrients contents. In [13], the authors grew *Arabidopsis* for 5 days at 16$h$ photoperiod in no-limiting nitrogen conditions ($n_{\text{soil}} = 12\mu molN/cm^3$) and phosphorus soil content equal to $p_{\text{soil}} = 0.125\mu molP/cm^3$. Then, plants were moved, for other 7 days, in a limiting soil ($p_{\text{soil}} = P_0 = 0\mu molP/cm^3$), a free soil ($p_{\text{soil}} = P_{0.125} = 0.125\mu molP/cm^3$) or toxic soils ($p_{\text{soil}} = P_{0.25} = 0.25\mu molP/cm^3$, $p_{\text{soil}} = P_{0.5} = 0.5\mu molP/cm^3$, $p_{\text{soil}} = P_1 = 1\mu molP/cm^3$ and $p_{\text{soil}} = P_2 = 2\mu molP/cm^3$). The authors found that:

- the shoot to root ratio is similar in the treatments $P_0$ and $P_{0.125}$, even if the total root length in the treatment $P_0$ is up to the 65% shorter than the total root length in $P_{0.125}$.
- the shoot to root ratio in the treatment $P_{0.125}$ is about two-fold the shoot to root ratio in $P_{0.25}$, while the total root length in $P_{0.25}$ is up to the 30% longer than in $P_{0.125}$;
- the shoot to root ratio is similar among the treatments $P_{0.5}$ and $P_1$ and $[10, 40]\%$ lower than the shoot to root ratio in $P_{0.125}$. The total root length in $P_{0.5}$ and $P_1$ is the $[0, 30]\%$ shorter than the $P_{0.125}$;
- the shoot to root ratio is the $[15, 45]\%$ higher when passing from the treatment $P_{0.125}$ to the treatment $P_2$, while the total root length decreases up to the 50%.

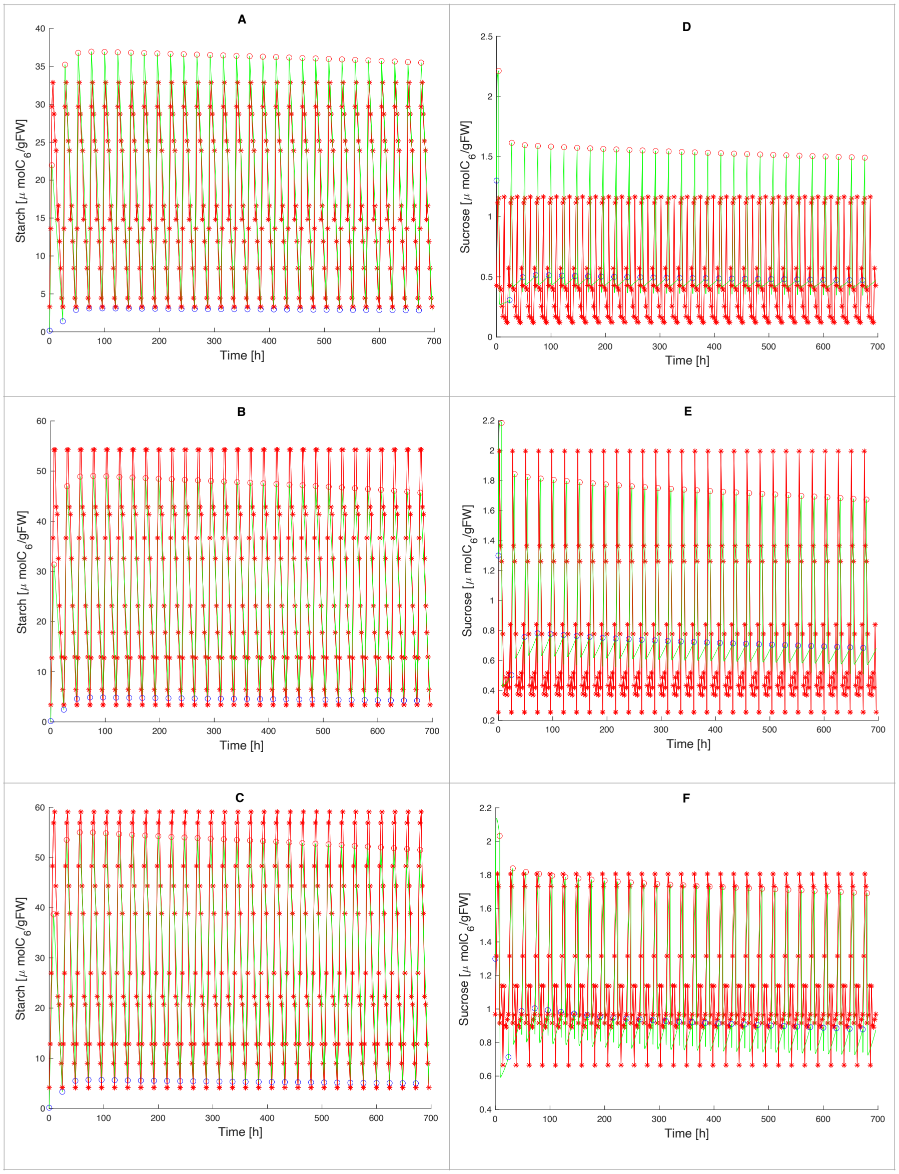


Figure 1: **Sucrose and starch evolution in plant growing with short photo-periods**. Red stars are data from [21]. In green, values from simulations. Blue and red dots indicate the dawn and the dusk, respectively.Starch (from A to C) and sucrose (from D to F) dynamics for $4h$ day length (in A and D), $6h$ day length (in B and E), $8h$ day length (in C and F).

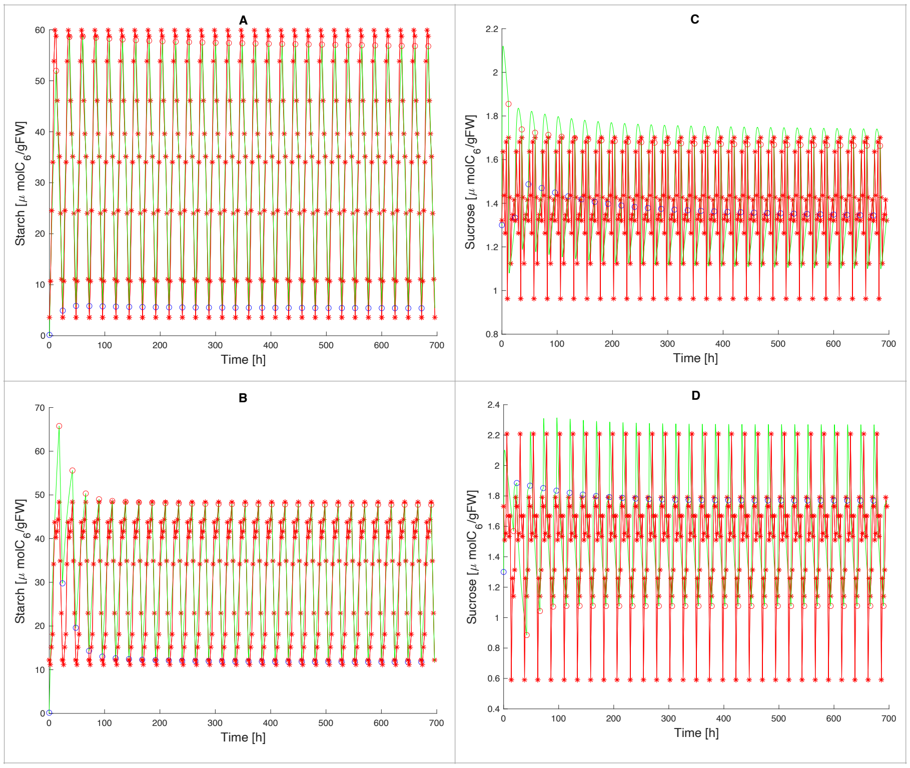


Figure 2: **Sucrose and starch evolution in plant growing with long photo-period**. Red stars are data from [21]. In green, values from simulations. Blue and red dots indicate the dawn and the dusk, respectively. Starch (in A and B) and sucrose (in C and D) dynamics for 12$h$ day length (in A and C), 8$h$ day length (in B and D).

We simulate 5 days of growth at 16$h$ photoperiod at $p_{\text{soil}} = 0.125$. Then $p_{\text{soil}}$ is changed according to the treatments $P_0$, $P_{0.125}$, $P_{0.25}$, $P_{0.5}$, $P_1$, $P_2$ and other 7 days of growth are simulated. In [13], authors measure the total root length. Here, we use the total root biomass to compare the results. Both the root biomass and the shoot to root ratio are reported in Table **??**.

In Table 2 we report the the total root biomass and the shoot to root ratio percentage variation (of limit and toxic conditions with respect to the control treatment $P_{0.125}$).

Let us note that almost all the results are in agreement with the observations

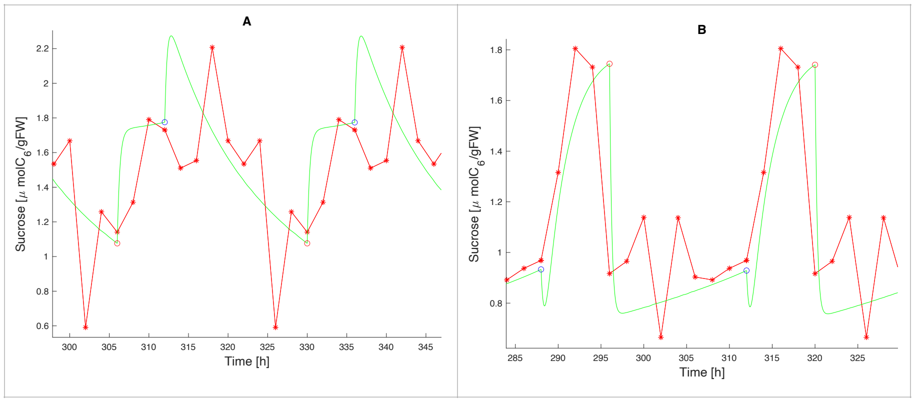


Figure 3: **Behaviour inaccuracies**. Red stars are data from [21]. In green, values from simulations. Blue and red dots indicate the dawn and the dusk, respectively. (A) Detail of sucrose content for $18h$ day length. (B) Detail of sucrose content for $8h$ day length.

Table 1: Total root biomass and shoot to root ratio.

| Treatment | Total root biomass $[\mu gFW]$ | Shoot to root ratio $[-]$ |
|---|---|---|
| $P_0$ | 25.14 | 1.36 |
| $P_{0.125}$ | 36.32 | 3.327 |
| $P_{0.25}$ | 47.43 | 1.83 |
| $P_{0.5}$ | 35.72 | 2.34 |
| $P_1$ | 31.28 | 1.934 |
| $P_2$ | 18.86 | 4.186 |

in [13]. Only the treatment $P_0$ underestimate the results. The reason could be that, to obtain the treatment $P_0$, the authors have replaced phosphorus with other elements. As noted in [13], the replacement could induce negative effects because of the toxic presence of other nutrients (that are not described into the model). The results in Table 1 have been used to estimate $\mu_1$, $\mu_2$, $\mu_5$, $\mu_6$ and

Table 2: Total root biomass and shoot to root ratio percentage variation with respect to $P_{0.125}$.

| Treatment | Total root biomass variation (%) | Shoot to root variation (%) |
|---|---|---|
| $P_0$ | -30.78 | -59.12 |
| $P_{0.25}$ | +30.58 | -44.99 |
| $P_{0.5}$ | -1.65 | -29.66 |
| $P_1$ | -13.87 | -41.86 |
| $P_2$ | -48 | +25.81 |

the following values for the function $p_{lux}(p_{\text{soil}})$:

$$p_{\text{lux}}(p_{\text{soil}}) = \begin{cases} 1 \quad p_{\text{soil}} \leq 0.125 \ \mu\frac{molP}{cm^3} \\ 1.6 \quad p_{\text{soil}} = 0.25 \ \mu\frac{molP}{cm^3} \\ 0.9 \quad p_{\text{soil}} = 0.5 \ \mu\frac{molP}{cm^3} \\ 1.3 \quad p_{\text{soil}} = 1 \ \mu\frac{molP}{cm^3} \\ 0.4 \quad p_{\text{soil}} = 2 \ \mu\frac{molP}{cm^3} \end{cases} .$$

The values of $p_{\text{lux}}$ when $p_{\text{soil}}$ is not in the previous cases are obtained by a spline interpolation.

Finally, in [6], the effects of poor and toxic nitrogen soil contents are estimated. The authors grew *Arabidopsis* for 35 days in $8h$ photoperiod in no-limiting phosphorus soil conditions ($p_{\text{soil}} = 0.15\mu molP/cm^3$). The nitrogen soil content is both very limiting ($n_{\text{soil}} = 0.89\mu molN/cm^3$), limiting ($n_{\text{soil}} = 7.5\mu molN/cm^3$), no-limiting ($n_{soil=15\mu molN/cm^3}$) and toxic ($n_{\text{soil}} = 30\mu molN/cm^3$). The shoot biomass is measured and compared with the shoot biomass of plants grown in Stender control soil ($n_{\text{soil}} = 20\mu molN/cm^3$). The authors observed that the highest values for the shoot biomass are obtained in the Stender control soil. Similar values are measured both in no-limiting and toxic conditions (the shoot biomass is $[0, 30]\%$ lower than the Stender control soil). In particular, the shoot biomass measured in the Stender control soil is $[60, 70]\%$ higher than the very limiting soil condition. In particular, according to this value we fix $\delta_{npd} = 0.3$). Finally, when passing from limiting to no-limiting soil condition the shoot biomass is $[20, 30]\%$ higher. When increasing the toxic levels of nitrogen soil content, the growth is reduced up to 30%. Then we fix $\delta_{nt} = 0.7$. In Table 3 we compare our results with experimental observations. The results are obtained by simulating 35 days of growth at $8h$ photoperiod with $p_{\text{soil}} = 0.15\mu molP/cm^3$ and $n_{\text{soil}} = \{0.89,\ 7.5,\ 15,\ 20,\ 30\}\mu molN/cm^3$. The initial biomass $b_l(0) = 6e-4gFW$ and $b_r(0) = 3e-4gFW$ are chosen so that the shoot biomass at the $35th$ day ($b_l(35)$) in the no-limiting condition ($n_{\text{soil}} = 15\mu molN/cm^3$) is in agreement with the experimental observations. The parameters $\mu_3,\ \mu_4,\ \mu_7,\ \mu_8,\ \mu_9,\ \mu_{10}$ are estimated to obtain the best fitting with the experimental observations.

Table 3: Total shoot biomass in $mgFW$.

| Treatment | Experimental range | Simulation |
|---|---|---|
| Very limiting | [60,100] | 84.98 |
| Limiting | [150,200] | 197.66 |
| No limiting | [220,300] | 262.63 |
| Stender control | [270,330] | 304.6 |
| Toxic | [220,300] | 268.16 |

### 3. LIST OF PARAMETERS

Table 4: List of Parameters

| Parameter | Value | Source | Significance |
|---|---|---|---|
| $s^{min}$ | 1.3 $[\frac{\mu mol C_6}{gFW}]$ | [23] | Lower bound sucrose content |
| $s^{max}$ | 2 $[\frac{\mu mol C_6}{gFW}]$ | [21] | Upper bound sucrose content |
| $p_h^{max}$ | 12.7 $[\frac{\mu mol C_6}{gFW \cdot h}]$ | [23] | Maximum photosynthetic rate |
| $\tau^{max}$ | 6 $[\frac{\mu mol C_6}{gFW \cdot h}]$ | [23] | Maximum starch degradation rate |
| $\delta_{pt}$ | 0.5 $[-]$ | [3] | Maximum reduction of $p_h^{max}$ because of toxic phosphorus levels |
| $n_{ph}$ | 5.75 $[\frac{\mu mol N}{gFW}]$ | [1] and equation (4) | Minimum nitrogen content to start photosynthesis |
| $a^{min}$ | 0.15 $[\frac{\mu mol C_6}{gFW}]$ | [24] | Minimum amount of starch at dawn |
| $p_c$ | 0.29 $[\frac{\mu mol C_6}{\mu mol P}]$ | [8] | Sugar cost of uptake phosphorus |
| $n_c$ | 0.65 $[\frac{\mu mol C_6}{\mu mol N}]$ | equation (24) | Sugar cost of uptake nitrogen |
| $\bar{r}_t$ | 0.0035 $[-]$ | [8] | Grams of carbon consumed for each gram of carbon loaded into the phloem |

Continued on next page

Table 4: List of Parameters (Continued)

| Parameter | Value | Source | Significance |
|---|---|---|---|
| $r_m^0$ | 0.79 $[\frac{1}{h}]$ | [25] | Frequency of sucrose loading into the phloem for the maintenance respiration |
| $r_g^0$ | 1.98 $[\frac{1}{h}]$ | [25] | Frequency of sucrose loading into the phloem for the growth respiration |
| $c_{sn}$ | 1.724 $[\frac{\mu molN}{\mu molC_6}]$ | [1] | Nitrogen consumed for each mole of carbon consumed during the respiration |
| $c_{sp}$ | 0.1724 $[\frac{\mu molP}{\mu molC_6}]$ | equation (21) | Phosphorus consumed for each mole of carbon consumed during the respiration |
| $\delta_{npd}$ | 0.3 $[-]$ | [6] | Minimum growth stimulus reduction due to soil nutrients deficiency |
| $\delta_{nt}$ | 0.7 $[-]$ | [6] | Minimum growth stimulus reduction due to soil nitrogen toxicity |
| $\bar{\mathcal{O}}$ | 10 $[\frac{\mu molN}{\mu molP}]$ | [2] | Optimal stoichiometry ratio in no-limiting conditions |

Continued on next page

Table 4: List of Parameters (Continued)

| Parameter | Value | Source | Significance |
|---|---|---|---|
| $\mathcal{O}^{min}$ | 3 $[\frac{\mu molN}{\mu molP}]$ | [14] | Minimum stoichiometry ratio |
| $\mathcal{O}^{max}$ | 30 $[\frac{\mu molN}{\mu molP}]$ | [14] | Maximum stoichiometry ratio |
| $I_n^{max}$ | 6.44 $[\frac{\mu molN}{gFW \cdot h}]$ | [18] | Maximum nitrogen uptake rate |
| $k_n^{max}$ | 0.125 $[\frac{\mu molN}{cm^3}]$ | [18] | Michaelis-Menten parameter for nitrogen uptake |
| $I_p^{max}$ | 0.494 $[\frac{\mu molP}{gFW \cdot h}]$ | [17] | Maximum phosphorus uptake rate |
| $k_p^{max}$ | 0.0074 $[\frac{\mu molP}{cm^3}]$ | [17] | Michaelis-Menten parameter for phosphorus uptake |
| $D$ | 4 $[-]$ | [20] | Memory days for thresholds computations |

| Parameter | Value | Source | Significance |
|---|---|---|---|
| $\lambda_{sdr}$ | 0.25 $[\frac{1}{h}]$ | calibration on [21] | Frequency parameter in $\gamma(t)$ dynamics (equation (12)) |
| $\lambda_{sdi}$ | 0.1 $[\frac{1}{h}]$ | calibration on [21] | Frequency parameter in $\gamma(t)$ dynamics (equation (12)) |
| $\lambda_{sni}$ | see below | calibration on [21] | Frequency parameter in $\gamma(t)$ dynamics (equation (12)) |
| $\lambda_c$ | see below | calibration on [21] | Control feedback on photosynthesis (equation (9)) |
| $\lambda_g$ | 0.65 $[-]$ | calibration on [21] | Control feedback on growth respiration (equation (22)) |
| $\lambda_{sb}$ | see below | calibration on [21] | Rate of sucrose conversion in new biomass |
| $\lambda_t$ | 1 $[h]$ | calibration on [21] | Frequency parameter in the equation (34) |
| $\lambda_{f1}$ | 0.023 $[\frac{\mu molN}{\mu molC_6}]$ | calibration on [21] | Conversion parameter in the equation (37) |
| $\lambda_{f2}$ | 0.01493 $[\frac{\mu molN}{\mu molC_6}]$ | calibration on [21] | Conversion parameter in the equation (37) |
| $\lambda_k$ | 50 $[-]$ | calibration on [21] | Weight of stoichiometry ration in the equations (45) |
| $\lambda_O$ | 0.0125 $[\frac{\mu molN}{\mu molP}]$ | calibration on [21] | Proportional parameter in the equation (28) |

Table 5: Dynamical model's variables.

| Variable | Unit of measure | Meaning |
|---|---|---|
| $t$ | $[s]$ | Time |
| $a(t)$ | $[\frac{\mu molC_6}{gFW}]$ | Starch content |
| $s(t)$ | $[\frac{\mu molC_6}{gFW}]$ | Sucrose content |
| $n(t)$ | $[\frac{\mu molN}{gFW}]$ | Nitrogen content |
| $p(t)$ | $[\frac{\mu molP}{gFW}]$ | Phosphorus content |
| $b_l(t)$ | $[gFW]$ | Leaf fresh biomass |
| $b_r(t)$ | $[gFW]$ | Root fresh biomass |
| $\gamma(t)$ | $[-]$ | Starch partition signal |
| $f_r(t)$ | $[-]$ | Root priority signal |
| $a_n(t)$ | $[-]$ | Nitrogen affinity signal |
| $a_p(t)$ | $[-]$ | Phosphorus affinity signal |

| Parameter | Value | Source | Significance |
|---|---|---|---|
| $\mu_1$ | $1.85$ $[-]$ | calibration on [13] | Parameter for toxic soil phosphorus effects (equation (8)) |
| $\mu_2$ | $-3$ $[\frac{cm^3}{\mu molP}]$ | calibration on [13] | Parameter for toxic soil phosphorus effects (equation (8)) |
| $\mu_5$ | $0.005$ $[-]$ | calibration on [13] | Parameter for limiting soil phosphorus effects (equation (18)) |
| $\mu_6$ | $56$ $[\frac{cm^3}{\mu molP}]$ | calibration on [13] | Parameter for limiting soil phosphorus effects (equation (18)) |
| $\mu_3$ | $0.6484$ $[-]$ | calibration on [6] | Parameter for limiting soil nitrogen effects (equation (18)) |
| $\mu_4$ | $0.0537$ $[\frac{cm^3}{\mu molN}]$ | calibration on [6] | Parameter for limiting soil nitrogen effects (equation (18)) |
| $\mu_7$ | $3.431$ $[-]$ | calibration on [6] | Parameter for toxic soil nitrogen effects (equation (18)) |
| $\mu_8$ | $-0.0642$ $[\frac{cm^3}{\mu molN}]$ | calibration on [6] | Parameter for toxic soil nitrogen effects (equation (18)) |
| $\mu_9$ | $1.842$ $[-]$ | calibration on [6] | Parameter for toxic soil nitrogen effects (equation (30)) |
| $\mu_{10}$ | $-0.0509$ $[\frac{cm^3}{\mu molN}]$ | calibration on [6] | Parameter for toxic soil nitrogen effects (equation (30)) |

| Parameter | Value | Source | Significance |
|---|---|---|---|
| $\theta_{bl}$ | $10^{-8}$ $[gFW]$ | | Shoot biomass parameter in the equation (1) |
| $\theta_w$ | $10^{-3}$ $[\frac{\mu molWater}{cm^3}]$ | | Water parameter in the equation (3) |
| $\theta_s$ | $0.005$ $[\frac{1}{h}]$ | | Rate of sucrose losses during respiration (17) |
| $\theta_{w2}$ | $1$ $[-]$ | | Water toxic effects on death rate of tissues |
| $\theta_{br}$ | $10^{-6}$ $[gFW]$ | | Root biomass parameter in the equation (35) |
| $\theta_{ld}$ | $5 \cdot 10^{-6}$ $[\frac{1}{h}]$ | | Death rate of leaves tissues |
| $\theta_{lc}$ | $5 \cdot 10^{-6}$ $[\frac{1}{h}]$ | | Competition rate of leaf biomass |
| $\theta_{rd}$ | $5 \cdot 10^{-6}$ $[\frac{1}{h}]$ | | Death rate of roots tissues |
| $\theta_{rc}$ | $5 \cdot 10^{-6}$ $[\frac{1}{h}]$ | | Competition rate of root biomass |

Table 6: Some parameters depend on the photoperiod. We estimated them for each day-length in [21]:

| | $4h$ | $6h$ | $8h$ | $12h$ | $18h$ |
|---|---|---|---|---|---|
| $\lambda_c$ $[-]$ | 0.82 | 0.79 | 0.67 | 0.62 | 0.55 |
| $\lambda_{sni}$ $[\frac{1}{h}]$ | 0.16 | 0.15 | 0.13 | 0.08 | 0.004 |
| $\lambda_{sb}$ $[\frac{gFW}{\mu molC_6}]$ | 0.002973 | 0.003493 | 0.004276 | 0.00475 | 0.005165 |

### 4. SUMMARY

The main equations characterising the dynamics are summarised below, while the main variables of the model are collected in the table 5.

*PHOTOSYNTHESIS*

$$p_h(t) = p_h^{max} L(t)\bar{w}(t)\min(\bar{n}(t), \bar{p}(t))p_{tox}(t, p_{soil}(t))C(t, a(t))\frac{b_l(t)}{b_l(t) + \theta_{bl}}.$$

*STARCH AND SUCROSE DYNAMICS*

$$\frac{da(t)}{dt} = \gamma(t)p_h(t) - \tau_{as}(t),$$
$$\frac{ds(t)}{dt} = (1 - \gamma(t))\,p_h(t) + \tau_{as}(t) - r^u(t) - r^t(t) - r^m(t) - r^g(t),$$
$$\frac{d\gamma(t)}{dt} = L(t)\left(-\gamma(t)\lambda_{sdr}\frac{s^{min}}{s(t) + s^{min}} + (1 - \gamma(t))\lambda_{sdi}\frac{s(t)}{s(t) + s^{max}}\right) + (1 - L(t))(1 - \gamma(t))\lambda_{sni}\frac{s^{min}}{s^{min} + s(t)}.$$

*MICHAELIS-MENTEN KINETICS*

$$u_{nMM}(I_n, k_n) = I_n\frac{n_{soil}}{n_{soil} + k_n},$$
$$u_{pMM}(I_p, k_p) = I_p\frac{p_{soil}}{p_{soil} + k_p}.$$

*NUTRIENTS UPTAKE*

$$
\begin{aligned}
u_n(t) &= u_{nMM}(I_n^{max}, k_n^{max}) a_n(t) u_n^{sat}(t) \frac{b_r(t)}{b_l(t) + b_r(t)}, \\
u_p(t) &= u_{pMM}(I_p^{max}, k_p^{max}) a_p(t) u_p^{sat}(t) \frac{b_r(t)}{b_l(t) + b_r(t)}, \\
\frac{da_n(t)}{dt} &= (1 - a_n(t)) \left( \left( 1 - \frac{u_n(t)}{u_n(t) + c_n(t) + \varepsilon} \right) \frac{n^{max}(t)}{n^{max}(t) + n(t)} + \frac{p_h(t)}{p_h^{max}} - a_n(t) \lambda_k (1 - 2 f_n(t)) \right) - \\
&\quad - a_n(t) \left( \frac{n(t)}{n(t) + n^{min}} + \frac{n_c u_n(t)}{n_c u_n(t) + p_h(t)(1 - \gamma(t)) + \tau_{as}(t) + \varepsilon} \right), \\
\frac{da_p(t)}{dt} &= (1 - a_p(t)) \left( \left( 1 - \frac{u_p(t)}{u_p(t) + c_p(t) + \varepsilon} \right) \frac{p^{max}(t)}{p^{max}(t) + p(t)} + \frac{p_h(t)}{p_h^{max}} + a_p(t) \lambda_k (1 - 2 f_n(t)) \right) - \\
&\quad - a_p(t) \left( \frac{p(t)}{p(t) + p^{min}} + \frac{p_c u_p(t)}{p_c u_p(t) + p_h(t)(1 - \gamma(t)) + \tau_{as}(t) + \varepsilon} \right).
\end{aligned}
$$

*NUTRIENTS DYNAMICS*

$$
\begin{aligned}
\frac{dn(t)}{dt} &= u_n(t) - c_n(t), \\
\frac{dp(t)}{dt} &= u_p(t) - c_p(t).
\end{aligned}
$$

*GROWTH AND SUCROSE ALLOCATION*

$$
\begin{aligned}
\frac{db_l}{dt} &= \lambda_{sb}(1 - f_r(t) n_{tox}(t)) r^g(t) b_l(t) - \theta_{w2}(t) \theta_{ld} b_l(t) - \theta_{lc} b_l^2(t), \\
\frac{db_r}{dt} &= \lambda_{sb} f_r(t) n_{tox}(t) r^g(t) b_l(t) - \theta_{rd} b_r(t) - \theta_{rc} b_r^2(t), \\
\frac{df_r(t)}{dt} &= p_{lux}(t)(1 - f_r(t)) \left( a_n(t) f_n(t) + (1 - f_n(t)) a_p(t) \right) - \\
&\quad - f_r(t) \left( \frac{n(t)(1 - f_n(t))}{n(t) + n^{min}(t)} + \frac{p(t) f_n(t)}{p(t) + p^{min}(t)} + \frac{s^{min}}{s(t) + s^{min}} \right).
\end{aligned}
$$
